# Proximity-labeling chemoproteomics defines the subcellular cysteinome and inflammation-responsive mitochondrial redoxome

**DOI:** 10.1101/2023.01.22.525042

**Authors:** Tianyang Yan, Ashley R. Julio, Miranda Villanueva, Anthony E. Jones, Andréa B. Ball, Lisa M. Boatner, Alexandra C. Turmon, Stephanie L. Yen, Heta S. Desai, Ajit S. Divakaruni, Keriann M. Backus

## Abstract

Proteinaceous cysteines function as essential sensors of cellular redox state. Consequently, defining the cysteine redoxome is a key challenge for functional proteomic studies. While proteome-wide inventories of cysteine oxidation state are readily achieved using established, widely adopted proteomic methods such as OxiCat, Biotin Switch, and SP3-Rox, they typically assay bulk proteomes and therefore fail to capture protein localization-dependent oxidative modifications. To obviate requirements for laborious biochemical fractionation, here, we develop and apply an unprecedented two step cysteine capture method to establish the Local Cysteine Capture (Cys-LoC), and Local Cysteine Oxidation (Cys-LOx) methods, which together yield compartment-specific cysteine capture and quantitation of cysteine oxidation state. Benchmarking of the Cys-LoC method across a panel of subcellular compartments revealed more than 3,500 cysteines not previously captured by whole cell proteomic analysis. Application of the Cys-LOx method to LPS stimulated murine immortalized bone marrow-derived macrophages (iBMDM), revealed previously unidentified mitochondria-specific inflammation-induced cysteine oxidative modifications including those associated with oxidative phosphorylation. These findings shed light on post-translational mechanisms regulating mitochondrial function during the cellular innate immune response.

## Introduction

Distinguished by their sensitivity to oxidative stress, proteinaceous cysteine residues play important roles in physiological processes and diseases, such as neurological disorders, cancers, and autoimmune disorders^1–4^. Interestingly, abnormal levels of reactive oxygen and nitrogen species (ROS and RNS) have been implicated in these diseases^5–8^. Cysteine chemoproteomic methods, such as biotin-switch^9^, Oxicat^10^, SP3-Rox^11^, and Oximouse^12^, enable high throughput quantitation of changes to cysteine oxidation states. Application of these methods have pinpointed cysteines differentially oxidized in association with high levels of ROS and RNS, such as those of TRX^13,14^, GAPDH^15,16^ and HBB^17^. Given the recent advent of cysteine-reactive small molecules as precision therapies for the treatments of cancers and immune disorders^18–20^, cysteine chemoproteomic methods have also emerged as enabling technology for pinpointing ligandable or potentially ‘druggable’ residues proteome-wide^21–30^. A central remaining challenge for these studies is the lack of a priori knowledge about the functional impact of covalent modification. Given the functional importance of cysteine oxidative modifications, understanding which cysteines serve as endogenous redox sensors is also of high utility for target prioritization efforts.

Nearly all cysteine redox profiling platforms follow the same general workflow: First, cells are lysed, and the reduced cysteines are capped with a pan-cysteine reactive reagent, such as iodoacetamide alkyne (IAA). After reduction, natively oxidized cysteines are then capped by another cysteine capping reagent, such as isotopically differentiated IAA. Samples are then biotinylated, enriched on avidin resin, subjected to sequence specific proteolysis, and liquid chromatography tandem mass spectrometry analysis (LC-MS/MS). Both absolute and relative changes to cysteine oxidation can be quantified either using MS1^21,22^ or MS2^12,31^-based quantification. While such studies provide a global snapshot of cysteine states, they fail to capture subcellular, compartment-specific changes in cysteine ligandability or oxidation state—such differences are to be expected given the established spectrum of organelle redox potentials^32,33^. Notably, by combining OxiCat with biochemical fractionation, recent studies have quantified cysteine oxidation for mitochondrial and endoplasmic reticulum localized proteins^34,35^. Such studies remain limited by the cumbersome nature of density gradient centrifugation, the propensity of cysteines to oxidize during sample manipulation, and incompatibility with membraneless organelles and other compartments for which subcellular fractionation is not feasible.

The emergence of proximity labeling techniques, including APEX^36^, BioID^37^, and TurboID^38^, has enabled high fidelity biotinylation and enrichment of proteins from a range of subcellular compartments, including the cytosol, nucleus, mitochondria, endoplasmic reticulum membrane and endoplasmic reticulum lumen. With the addition of exogenous biotin (or related biotin analogues), biotinylation occurs with spatiotemporal control inside the targeted organelle. Pioneering studies have demonstrated the utility of these proximity based labeling methods in deciphering the protein interactome^39,40^, the protein composition of membraneless organelles^41,42^, and interrogation of kinase substrates^43^. Whether these methods are compatible with capturing the subcellular redoxome remains to be seen.

Across all organelles, the mitochondrial cysteine redoxome is particularly intriguing. In addition to carrying out oxidative phosphorylation (OXPHOS), mitochondria also play key roles in nearly all aspects of cell physiology, including functioning as hubs for biosynthesis, Ca^2+^ handling, iron homeostasis, and signal transduction^44,45^. Additionally, mitochondria are also thought to be significant producers of reactive oxygen species (ROS)^46^, and ROS-sensitive cysteines are known to regulate mitochondrial proteins such as aconitase and respiratory complex^47–49^. Mitochondrial ROS is also an emerging hallmark of the innate immune response^50,51^. For example, in response to lipopolysaccharide (LPS), murine bone marrow-derived macrophages (BMDMs) adopt a pro-inflammatory program which includes the near-total collapse of mitochondrial oxidative phosphorylation^52–54^. LPS induces an inflammatory response by initiating a signaling cascade that leads to NFkB translocation and expression of proinflammatory cytokines^55^. ROS and RNS levels profoundly increase during inflammatory processes, in part due to expression of inducible nitric oxide synthase (iNOS)^56^. However, the extent to which specific mitochondrial cysteines are oxidized as a result of this mitochondrial reprogramming remains largely unknown.

Here we combine enzymatic (TurboID) proximity based biotinylation with cysteine redox state analysis to enable in situ subcellular cysteine fractionation and quantitative measures of cysteine oxidation state. We first established the Local Cysteine Capture (Cys-LoC) method, which, when applied to cells expressing TurboID localized to cytosol, endoplasmic reticulum (ER), mitochondria (Mito), golgi, and nucleus, identified >3,500 cysteines not previously captured by whole cell proteomic analysis^25^. On average, 500 cysteines were captured from each compartment that were not enriched from HEK293T whole cell lysates. Unexpectedly, we observed low subcellular specificity for constructs targeted to a subset of compartments, which we mitigated through simultaneous depletion of endogenous biotin and translation arrest-induced depletion of newly translated TurboID. By combining these two innovations with our SP3-Rox method^11^, we then established the Local Cysteine Oxidation (Cys-LOx) method. When applied to identify inflammation-sensitive cysteines in an immortalized bone marrow derived murine macrophage (iBMDM) cell line exposed to LPS, we identified 32 mitochondria specific cysteines that exhibited cell-state dependent oxidation, including residues in proteins important for respiration, associated with oxidative phosphorylation and those not captured using bulk SP3-Rox analysis.

## Results

### Establishing the Local Cysteine Capture (Cys-LoC) method accesses the subcellular cysteineome

Here we envisioned combining proximity labeling via the ultra-fast biotin ligase TurboID with cysteine chemoproteomics^11,21,24–30,57,58^ to enable fractionation-free capture of the subcellular cysteinome, for both residue identification and quantification of cysteine oxidation. We were inspired by recent reports of two-step capture for subcellular phosphoproteomics, in which proteins biotinylated by TurboID were first enriched on avidin resin followed by peptide-level capture of phosphopeptides^59^. As a first step to test the feasibility of an analogous two step enrichment method for cysteine chemoproteomics, we transiently overexpressed a panel of TurboID fusion proteins tagged with localization sequences targeted to cytosol (cyto), endoplasmic reticulum (ER), golgi, mitochondrial (mito), and nucleus (nuc) (**Figure S1**)^38,60^. We then combined expression of these constructs with a customized two-step enrichment strategy, termed Local Cysteine Capture (Cys-LoC) (**Figure 1A**). In Cys-LoC, TurboID proximal proteins are first biotinylated in situ. Following lysis and cysteine capping with the highly reactive iodoacetamide alkyne (IAA), biotinylated proteins are enriched on streptavidin resin and subjected to sequence specific proteolysis. This digest releases all IAA-tagged peptides derived from the TurboID modified proteins. Subsequent peptide-level click conjugation to biotin-azide followed by a second enrichment on neutravidin resin affords specific capture of biotinylated cysteine peptides derived from turboID-modified proteins. Demonstrating the utility of the Cys-Loc method, we found that coverage of cysteines substantially increased with two-step biotinylation based Cys-LoC (**Figure 1A**) compared to one-step TurboID (**Figure S2**).

**Figure 1.**
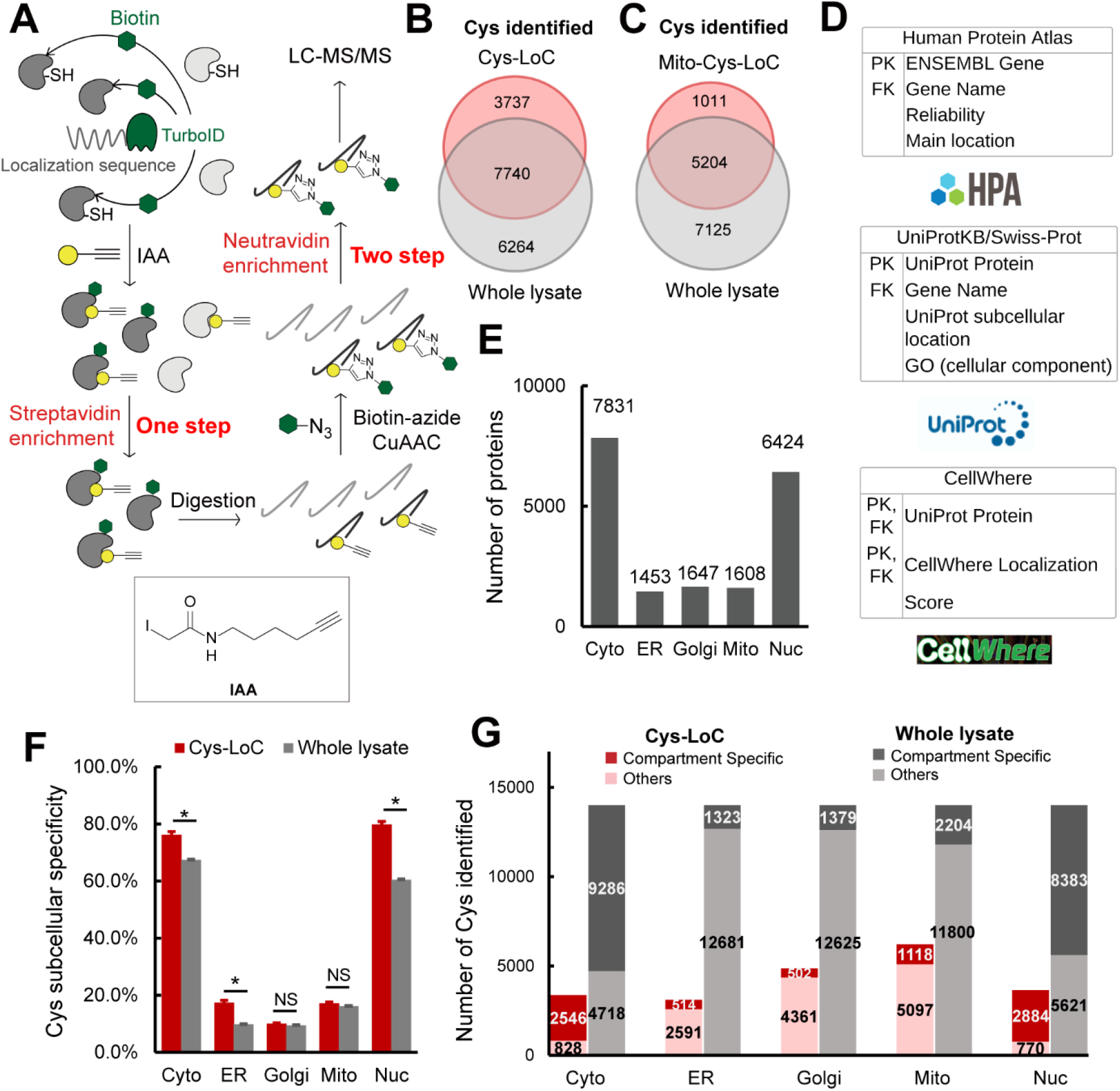
Establishment of Local Cysteine Capture (Cys-LoC) method. **A)** Scheme of Cys-LoC Workflow. **B)** Cysteines identified with Cys-LoC from five compartments aggregated compared to those identified from whole proteome in HEK293T cells^25^. **C)** Cysteines identified with Mito-Cys-LoC compared to those identified from the whole proteome in HEK293T cells^25^. **D)** Scheme of database generation with aggregated protein localization annotations from Human Protein Atlas, UniprotKB and CellWhere. PK and FK represented primary key and foreign key, respectively. **E)** Number of proteins annotated as localized in cytosol (cyto), endoplasmic reticulum (ER), golgi, mitochondrial (mito), and nucleus (nuc). **F)** Subcellular specificity of cysteines identified with Cys-LoC or with whole lysate [cite sp3 paper]. **G)** Number of cysteines identified as compartment specific with Cys-LoC or with whole lysate in HEK293T cells^25^. Statistical significance was calculated with unpaired Student’s t-tests, * p<0.05, NS p>0.05. Experiments were performed in duplicates in HEK293T cells for penal B, C, F, G. All MS data can be found in **Table S1**.

Implementation of the Cys-LoC method for constructs targeted to all five aforementioned compartments identified in aggregate 11,478 total cysteines, with an average of 3,700 cysteines per construct (**Figure 1B**). Gratifyingly, more than 450 cysteines were identified from each compartment that had not been previously captured in our previous bulk cysteinome analysis of HEK293T using out SP3-FAIMS method^25^ (**Figure S3**). Further exemplifying the utility of the Cys-LoC method to capture novel cysteines, for the mitochondrial targeted construct, 1,011 cysteines were identified that were not previously captured by our prior study^25^ (**Figure 1C**). When cross referenced with our newly reported CysDB database of 62,888 total identified cysteines, 489 cysteines identified by Cys-LoC had not been previously reported by any of our panel of high coverage cysteine chemoproteomics studies^24^.

### Evaluating the subcellular specificity of the cysteines captured by Cys-LoC

Motivated by the observed expanded portrait of the cysteinome enabled by Cys-LoC, we next asked whether the cysteine-containing proteins captured were representative of the subcellular compartments to which the respective TurboID constructs were targeted. To facilitate the analysis of subcellular proteomes, we generated a comprehensive protein localization database by aggregating protein localization information from the Human Protein Atlas^61^, UniprotKB^62^ and CellWhere^63^ (**Table S1**). Of the 16,983 proteins with available localization information, 12,835 human proteins were annotated as localized in the cytosol, ER, golgi, mitochondria and nucleus (**Figure 1D**). 7,831, 1,453, 1,647, 1,608, and 6,424 proteins were annotated as localized in the cytosol, ER, golgi, mitochondria and nucleus, respectively (**Figure 1E**).

Stratification of our Cys-LoC dataset by TurboID subcellular localization revealed several striking features. For the constructs targeted to the cytosol and nucleus, we observed comparatively high (~80%) localization specificity, calculated as the percentage of cysteines identified with protein localization annotations matching the compartment targeted by the respective TurboID (**Figure 1F**). However, when this analysis was extended to the ER, mito and golgi datasets, the specificity dropped dramatically (<20%). While some variability in compartment-specific proximity labeling has been reported previously^38^, the scale of the difference between compartments was unexpected.

We next interrogated how the compartment specificity achieved by Cys-Loc compared to datasets generated from unfractionated HEK293T whole cell proteome^24,25^. We observed modest, yet statistically significant, enrichment for cysteines captured using the Cyto-, Nuc- and ER-Cys-LoC platforms. In contrast, no significant enrichment was observed for the Golgi and Mito-targeted constructs (**Figure 1F**). Extension of this analysis to consider the total number of cysteines identified revealed a marked decrease in coverage for all constructs assayed via Cys-LoC when compared with whole lysate analysis. For example, cysteine chemoproteomics analysis of whole cell lysate identifies 2,000 mitochondria localized cysteines. The mito-Cys-LoC platform decreases the number of background cysteines (from 11,800 to 5,097) alongside the number of mitochondria cysteines (from 2,204 to 1,118) (**Figure 1G)**. Nonetheless, 3,737 cysteines were identified by the Cys-LoC platform that had not been identified in our prior SP3-FAIMS analysis of matched HEK293T proteome, including C588 of presequence protease (PITRM1)^64^, a protease responsible for clearance of accumulated mitochondrial amyloid beta protein, as well as zinc-coordinating cysteine C256 of DnaJ homolog subfamily A member 3 (DNAJ3), a protein regulator of apoptotic signaling in cancer^65^ (**Figure 1B, 1C, S3**). These examples highlight the utility of Cys-Loc enabled in situ subcellular fractionation for uncovering novel cysteines. More notable still, 489 cysteines had not been previously identified in any of the high coverage cysteine chemoproteomic datasets aggregated in our CysDB database (out of a total of 62,888 cysteines)^24^.

### Investigating established parameters associated with TurboID performance

For most proximity-labeling studies, some background labeling can be accommodated with appropriate controls (e.g. +-treatment groups). In contrast, implementation of proximity labeling to measure compartment specific changes to the cysteinome, for example for cysteine oxidation, requires comparatively high compartment labeling specificity, as rationalized by the following hypothetical cysteine: for a cysteine that is heavily oxidized in the mitochondria but not in the cytoplasm, proximity labeling that captures both subsets of the protein would incorrectly report the average oxidation state across both compartments. With the goal of minimizing non-specific cysteine enrichment, we sought to first pinpoint and then address sources of the observed seemingly promiscuous proximity labeling.

To streamline our efforts at method optimization, we established a protein-level proximity labeling workflow (**Figure 2A**) in which TurboID specificity was assayed by the fraction of total proteins identified in which the localization matched that of the TurboID fusion protein. Using this platform paired with transient overexpression we observed comparable protein localization specificity to that achieved with the Cys-LoC method (**Figure 2B and Figure 1E**), indicating that the peptide-level enrichment analysis in the Cys-LoC workflow was not a significant contributor to the low specificity.

**Figure 2.**
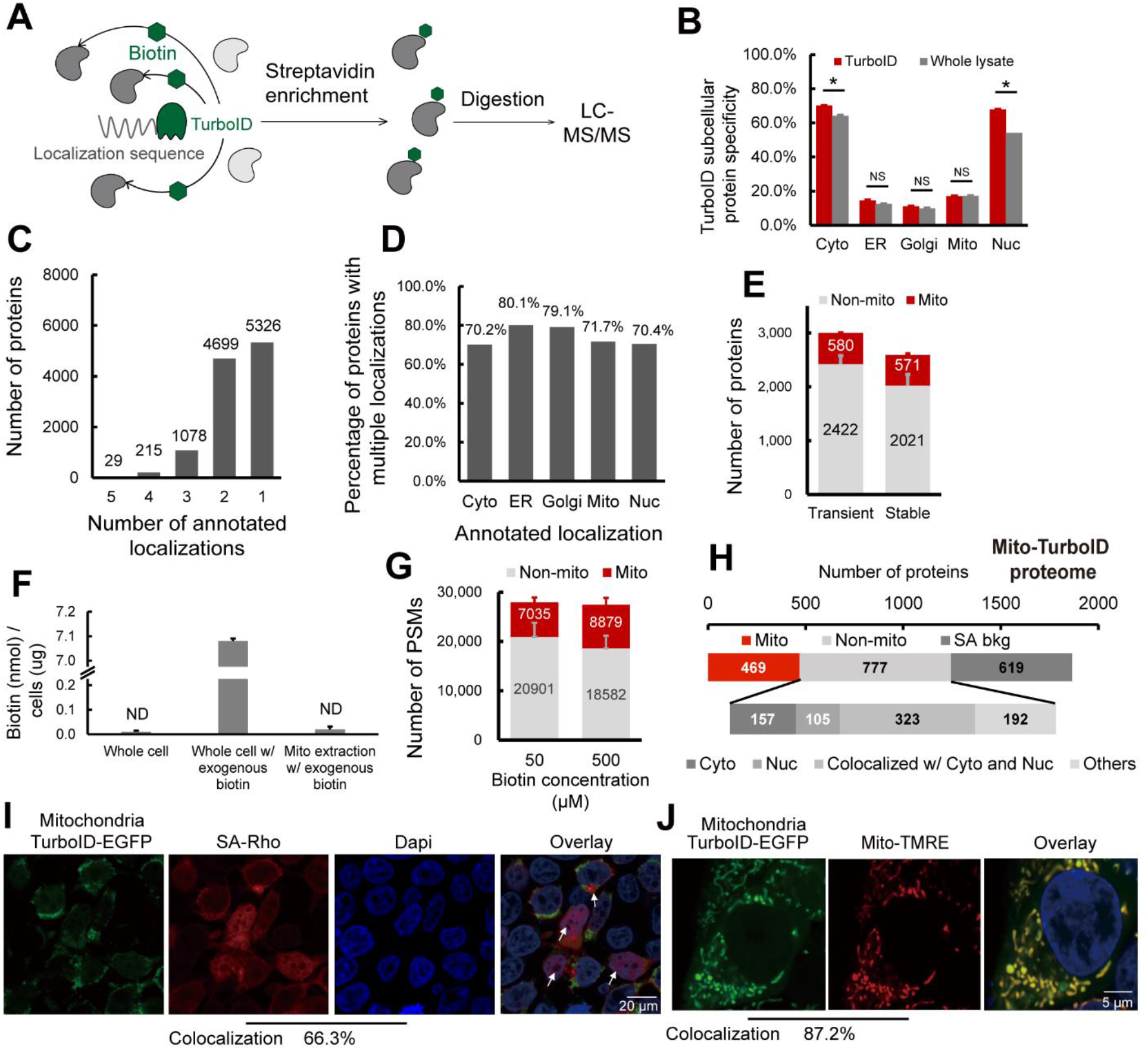
Pinpointing sources of non-compartment specific TurboID biotinylation. **A)** Scheme of TurboID based proximity labeling protein enrichment. **B)** Subcellular specificity of proteins identified with TurboID or with whole lysate. **C)** Number of proteins by number of localization annotations. **D)** Percentage of proteins within each subcellular compartment containing greater than 1 annotated localization. **E)** Number of mitochondrial annotated proteins and non-mitochondrial annotated proteins identified with transiently or stably expressed mito-TurboID. **F)** Absolute quantification of biotin detected for the whole cell, the whole cell with 500 μM exogenous biotin added, and the mitochondrial fraction with 500 μM exogenous biotin added. **G)** Number of PSMs of mitochondrial annotated proteins and non-mitochondrial annotated proteins identified with mito-TurboID with 50 μM or 500 μM exogenous biotin. **H)** Proteome analysis of the proteins enriched with mito-TurboID. SA-bkg indicated proteins identified in streptavidin background proteome. **I)** Localization of biotinylation in cells with mito-TurboID indicated by signals of streptavidin-rhodamine (SA-Rho). **J)** Localization of mito-TurboID overlaid with Mito-TMRE. Statistical significance was calculated with unpaired Student’s t-tests, * p<0.05, NS p>0.05. Experiments were performed in duplicates in HEK293T cells for panel B, E, F, G, H. All MS data can be found in **Table S2**.

We then investigated whether our protein localization database might provide insights into why some TurboID constructs show improved performance. Many proteins are known to have multiple subcellular localizations. Within our protein localization database, we found that there are more than 6,000 proteins with multi-localization annotations, including 29 proteins annotated as localized in all five compartments, such as cGMP-dependent 3’,5’-cyclic phosphodiesterase PDE2A, E3 ubiquitin-protein ligase parkin PRKN and the family of MAP kinase-activated protein kinases MAPKs, all of which have been reported to mediate a diverse array of functions across multiple subcellular locations^66–68^(**Figure 2C and Table S1**). Notably, ER-localized proteins showed the most pronounced multi-localization with 80% labeled are belonging to multiple compartments (**Figure 2D**). Removal of these multi-location proteins from our datasets substantially decreased performance of Cys-LoC (**Figure S4**). These findings further supported the necessity of a comprehensive protein localization dataset, such as we have presented in **Table S1**, which aggregates available protein localization data. Unsatisfied with the results of post-acquisition data filtration in improving Cys-Loc performance, we turned to our experimental workflow, seeking to increase specificity through methodological optimization.

As one of our primary goals in establishing the Cys-Loc method was to enable streamlined mitochondrial redox proteomics, we opted to use our mitochondrial targeted construct to perform further in-depth analysis of our protocol. Consistent with the previous study that achieved decreased non-specific biotinylation by regulating TurboID expression with an inducible construct^69^, we found that stable expression of the mito-TurboID compared to transient expression, afforded an increase in mitochondrial protein specificity (22% vs 19%) with only a modest decrease in net proteins identified (571 vs 580, **Figure 2E**).

Given that previous reports indicated comparatively low specificity for TurboID-catalyzed labeling of mitochondria^38^, we hypothesized that low local biotin concentration might contribute to decreased specificity for mitochondrial proteins. While mitochondrial biotin uptake has been suggested to occur through passive diffusion, the prior report of pH dependent uptake is suggestive of saturable mitochondrial biotin levels^70^. We find that, after a pulse with 500 μM exogenous biotin, the absolute detectable levels of biotin in the cytoplasm rises rapidly, reaching 7.08 nmol/μg cells. In contrast, biotin remained below the limit of detection in crude mitochondrial extracts (**Figure 2F** and **Figure S5**). While TurboID has been shown to proceed efficiently at low biotin concentrations (50 μM), due to its increased (relative to BioID) affinity for biotin^38,69^, our findings pointed towards the possible requirement for increased biotin concentrations to achieve efficient labeling of mitochondrial proteins. Supporting this premise, comparison of 50 μM and 500 μM biotin revealed increased peptide spectrum matches (PSMs) for mitochondrial proteins with elevated biotin concentrations (**Figure 2G**).

Given TurboID’s enhanced labeling kinetics relative to BioID^38^, we investigated labeling time and observed that an improved balance of coverage and specificity could be achieved with 1h labeling time, when compared to 10 min or 3h (**Figure S6A**). As decreased biotinylation was observed in higher passage (>10) cell lines (**Figure S6B**), we restricted subsequent analyses to cell lines with <10 passages.

Taken together, implementation of stable expression of the TurboID fusion protein, 1h labeling time and 500 μM biotin afforded a modest improvement of mito-TurboID specificity to 27% (449/1690).

### Promiscuous Mito-TurboID labeling of cytosolic and nuclear proteins

To further understand the factors contributing to our still marginal compartment specificity, we asked whether insights could be garnered by analyzing the annotated localization of enriched proteins. To eliminate the possibility of non-specific streptavidin binding confounding our analysis, we generated a dataset (**Table S2** and methods for details) that defines the HEK293T streptavidin background proteome. Excluding the streptavidin background, we find that >75% (585/777) of the non-mitochondrial proteins enriched by mito-TurboID are annotated as localized in the nucleus or cytosol (**Figure 2H**). Examples of these non-mitochondrial proteins captured by the mitochondrial-targeted TurboID include histones H2B, H1and H3, ribosomal protein S6 kinase RPS6KA3, eukaryotic translation initiation factors EIF3M, EIF2D and ELF6. All these proteins are closely related to protein translation, suggestive of proximity labeling by newly translated, not yet localized TurboID-fusion proteins.

Site-of-labeling studies enabled via capture and LC-MS/MS analysis of TurboID biotinylated peptides further confirmed these findings with 66% (4,934/7,449) of all captured peptides stemming from proteins with nuclear localization. By comparison only 14% (1,021/7,449) of biotinylated peptides were derived from mitochondrial proteins (**Figure S7**). This marked labeling of nuclear proteins was also visualized by immunocytochemistry (ICC) in which a substantial accumulation of streptavidin labeling was observed in cellular nuclei (66.3% colocalization with DAPI), indicated by the white arrows (**Figure 2I**). By comparison, the mitochondrial localization of the Mito-TurboID-EGFP was observed to be high, as indicated by 87.2 % colocalization with mitotracker-tetramethylrhodamine ethyl ester (TMRE) (**Figure 2J**).

### Newly translated TurboID is a major cause of promiscuous biotinylation, which can be improved through translation arrest and depletion of endogenous biotin

Inspired by the modest, yet detectable signal for mito-TurboID detected by ICC outside of the mitochondria (**Figure 2I**), in addition to the fact that many non-mitochondrial proteins captured by mito-TurboID were closely related to protein translation, we postulated that this trace signal might stem from newly translated protein. To test this hypothesis, we performed subcellular fractionation to quantify the fraction of mito-TurboID localized to mitochondrial versus non-mitochondrial compartments. 30% of the total mito-TurboID was found localized outside of the mitochondria (**Figure 3A**). The non-mitochondrial fraction of mito-TurboID protein was nearly completely eliminated in cells subjected to translation arrest (Cycloheximide^71^, 100 ug/mL, 6h; **Figure 3B, 3C**), which further implicates newly translated TurboID protein as the likely source of the observed low subcellular labeling specificity. Further supporting this model, we observed a comparatively long (>24h) half-life for the mito-TurboID protein (**Figure 3D**). Proteomic analysis of CHX-treated cells revealed an increase in the mitochondrial protein specificity to 30% of all proteins enriched (335/1120). Consistent with the CHX treatment primarily impacting newly translated rather than mitochondrial localized proteins, we only observed a modest decrease in total mitochondrial proteins identified (<10%), with a more pronounced decrease in non-mitochondrial proteins (>20%) (**Figure 3E**). The impact of cycloheximide treatment was also detectable by streptavidin blot, where several bands showed a pronounced decrease is staining upon cycloheximide treatment (**Figure S8, * bands**).

**Figure 3.**
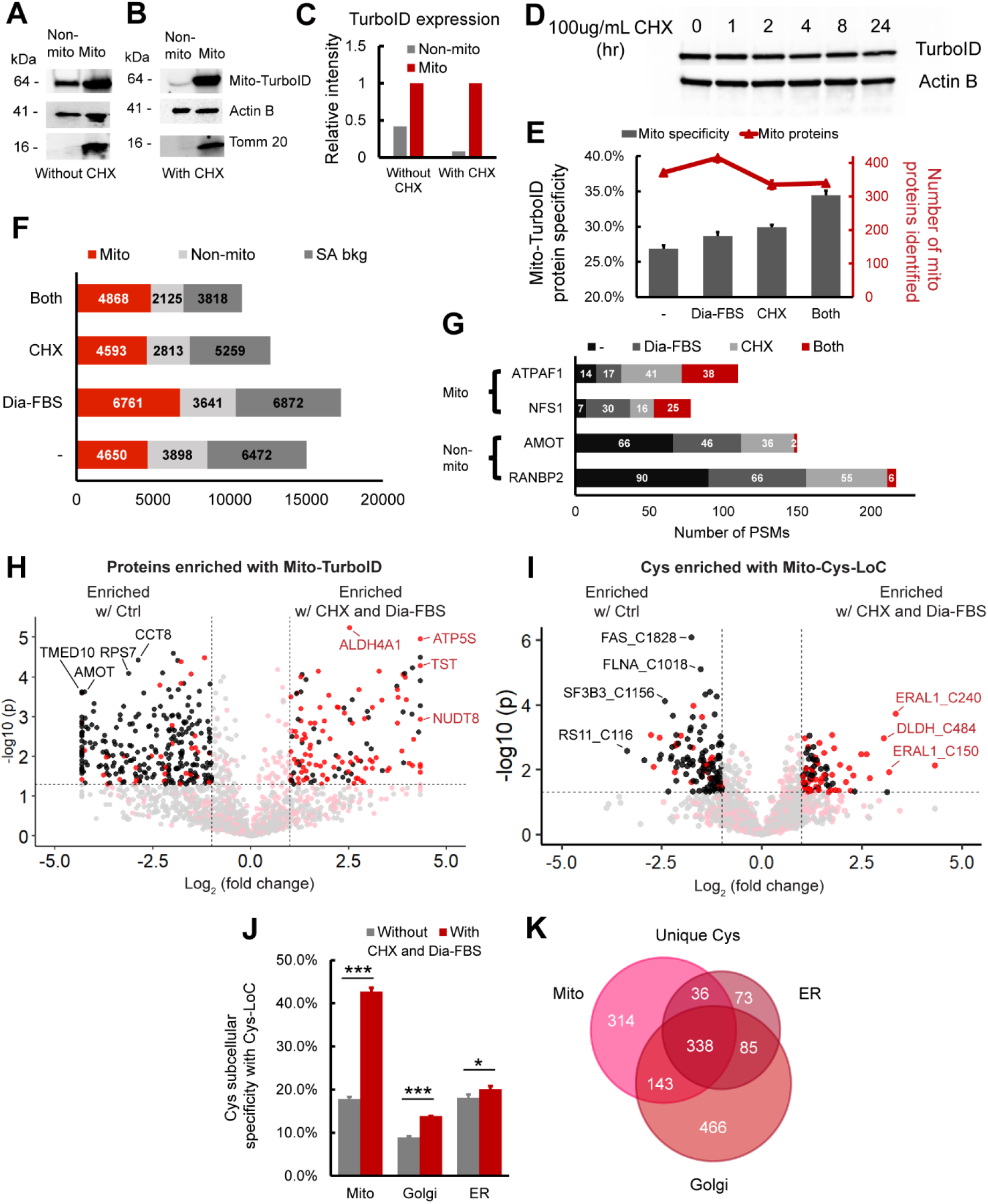
Translation arrest improves the subcellular specificity of proteins enriched by TurboID and cysteines captured by Cys-LoC. **A)** Abundance of mito-TurboID in mitochondrial vs non-mitochondrial fractions. Non-mitochondrial fraction refers to the rest of the lysate after crude mitochondria extraction. **B)** Abundance of mito-TurboID in mitochondrial vs non-mitochondrial fractions with CHX treatment. **C)** Quantitation of blot data in **3A** and **3B**. Intensity of the Mito-TurboID lane in the non-mitochondrial fraction is normalized to that in the mitochondrial fraction, which is set to 1.0. **D)** Expression of TurboID upon treatment of the translational inhibitor CHX. **E)** Specificity and number of mitochondria localized proteins enriched with mito-TurboID for control condition, dialyzed FBS, CHX treatment or for both. **F)** Distribution of the proteins enriched with mito-TurboID for control condition, dialyzed FBS, CHX treatment or both. **G)** Number of PSMs of representative mitochondrial proteins and non-mitochondrial proteins enriched with mito-TurboID for control condition, dialyzed FBS, CHX treatment or with both. **H)** Difference in intensities of proteins enriched with mito-TurboID with or without dialyzed FBS and CHX treatment. Red dots indicate proteins localized in mitochondria. Black dots indicate proteins localized in organelles other than mitochondria. **I)** Difference in intensities of cysteines enriched with Mito-Cys-LoC with or without dialyzed FBS and CHX treatment. Red dots indicate cysteines localized in mitochondria. Black dots indicate cysteines localized in organelles other than mitochondria. **J)** Specificity of cysteines identified in different subcellular compartments after Cys-LoC with or without dialyzed FBS and CHX treatment and with TurboID optimization. **K)** Cysteines identified with Cys-LoC targeting different subcellular compartments. Dialyzed FBS (Dia-FBS) treatment was 36 h and CHX treatment was 100 ug/mL for 6 h at 37 °C. Statistical significance was calculated with unpaired Student’s t-tests, * p<0.05, ** p<0.01, *** p<0.005. NS p>0.05. Experiments were performed in triplicates in HEK293T cells. All MS data can be found in **Table S3**.

Streptavidin blot analysis additionally indicated the presence of CHX-insensitive bands in biotin un-treated samples (**Figure S8**). Prior reports have implicated endogenous biotin as a substrate of TurboID and a source of background labeling; biotin-free dialyzed serum was recently found to decrease this low-level labeling^69,72^. Consistent with these findings, we observed that proximity labeling using dialyzed fetal bovine serum (Dia-FBS) afforded a modest but significant increase in mitochondrial protein specificity (27% to 29%) together with slight increase in overall mitochondrial proteins identified (415/1448; **Figure 3E**). More striking, when the dialyzed FBS and CHX treatments were combined nearly 35% of all proteins identified were mitochondrial, and no further decrease in protein coverage (340/987) was observed compared to CHX treatment alone. These findings were further substantiated by streptavidin blot visualization of decreased signal for CHX- and Dia-FBS-sensitive bands both in the presence and absence of exogenous biotin (**Figure S8, * bands**).

An increase from 27% to 35% mitochondrial protein specificity might, at first glance, appear relatively modest and inconsequential. Stratification of identified PSMs, rather than proteins, by annotated protein localization, more substantially revealed the impact of the combined CHX and Dia-FBS treatment. Excluding PSMs derived from background streptavidin binding, we find that the CHX and Dia-FBS treatment affords a 45% decrease in non-mitochondrial PSMs, from 3898 to 2125 PSMs. In contrast, the mitochondrial PSMs showed a modest increase from 4650 to 4868 with the CHX and Dia-FBS conditions (**Figure 3F**). Further exemplifying the impact of the combined translation arrest and biotin treatment, the number of PSMs for some of the most substantially enriched nuclear and cytoplasmic proteins, including angiomotin AMOT, a component of the 40S ribosomal subunit involved in translational repression^73^ and E3 SUMO-protein ligase RANBP2, which facilitates SUMO1 and SUMO2 conjugation^74^, decreased by 20-fold, whereas exemplary mitochondrial proteins ATP synthase mitochondrial F1 complex assembly factor 1 ATPAF1, which supports mitochondiral respiration^75^ and cysteine desulfurase NFS1, which catalyzes the desulfuration of L-cysteine to L-alanine^76^, remained unaffected by the treatment (**Figure 3G**). Label free quantification (LFQ) comparing relative abundance of proteins captured by TurboID in standard vs CHX-Dia-FBS treated samples revealed preferential enrichment of mitochondrial proteins with CHX-Dia-FBS (90/136 proteins enriched > 2-fold, red dots, **Figure 3H)** compared with preferential capture of non-mitochondrial proteins under normal treatment conditions (231/265 proteins enriched > 2-fold, black dots, **Figure 3H)**.

Extension of these analyses to Cys-LoC captured cysteine peptides confirmed that the dual CHX/Dia-FBS treatment afforded comparable increased performance to that observed for protein-level analysis. LFQ analysis revealed that 52% (47/90) of cysteines preferentially captured with CHX/Dia-FBS treatment belonged to mitochondrial localized proteins whereas nearly all (97/120) of those preferentially enriched under normal treatment conditions belonged to non-mitochondrial proteins (**Figure 3I**). Of note, only a handful of mitochondrial cysteines with more pronounced capture under normal conditions, including notably those found in cytochrome c oxidase subunit 4 isoform 1 COX4I1, which drives oxidative phosphorylation^77^ and receptor of activated protein C kinase 1 RACK1 proteins. These proteins are known to have a comparatively short half-life^78^. In aggregate, the CHX-Dia-FBS treatment, together with TurboID optimization mentioned beforehand, increased Cys-LoC mitochondrial cysteine specificity from 18% to 43% (**Figure 3J**).

As our methodological optimization was overwhelmingly focused on improving capture of mitochondrial cysteines, we next asked whether the improvement in specificity afforded by the CHX-Dia-FBS treatment would extend to other low specificity TurboID constructs. Cys-LoC analysis of HEK293T cells stably expressing either a Golgi- or ER-targeted TurboID construct revealed more modest but still significant increases in cysteine localization specificity (**Figure 3J**). The overlap in cysteines identified comparing the Mito-, Golgi- and ER-TurboID expressing cell lines was comparatively modest, spanning 55%-86% across three replicates analyzed (**Figure 3K**). Among the cysteines captured, several residues stood out for their lack of detection in prior studies (e.g. CysDB^24^), including C21 of oxygen-dependent coproporphyrinogen-III oxidase (CPOX), an enzyme that contributes to heme biosynthesis^79^, and C75 of DNL-type zinc finger protein (DNLZ), which coordinates a zinc in the zinc finger of this mitochondrial chaperone^80^.

### Establishing and applying the Local Cysteine Oxidation (Cys-LOx) method to analyze basal mitochondrial cysteine oxidation

Having achieved a substantial performance increase for Cys-LoC specificity, we returned to our second original objective, quantification of local cysteine oxidation state. To establish the Local Cysteine Oxidation (Cys-LOx) method (**Figure 4A**),proteins proximal to mitochondria were biotinylated via mito-TurboID. The cells were then lysed and reduced cysteines immediately capped using our custom isotopically labeled isopropyl iodoacetamide alkyne (IPIAA-L) probe^11^. Subsequently, the samples were subjected to reduction and capping using our isotopically differentiated heavy (IPIAA-H) probe. Fragpipe IonQuant^81^ was applied to report the ratio of IPIAA-L vs IPIAA-H labeled cysteine peptides (Log_2_(H/L)), from which the cysteine % oxidation state was calculated. As with Cys-LoC, the key innovative step of Cys-LOx is our unprecedented two-step enrichment protocol, which allows for capture of mitochondrial cysteines without conventional biochemical fractionation.

**Figure 4.**
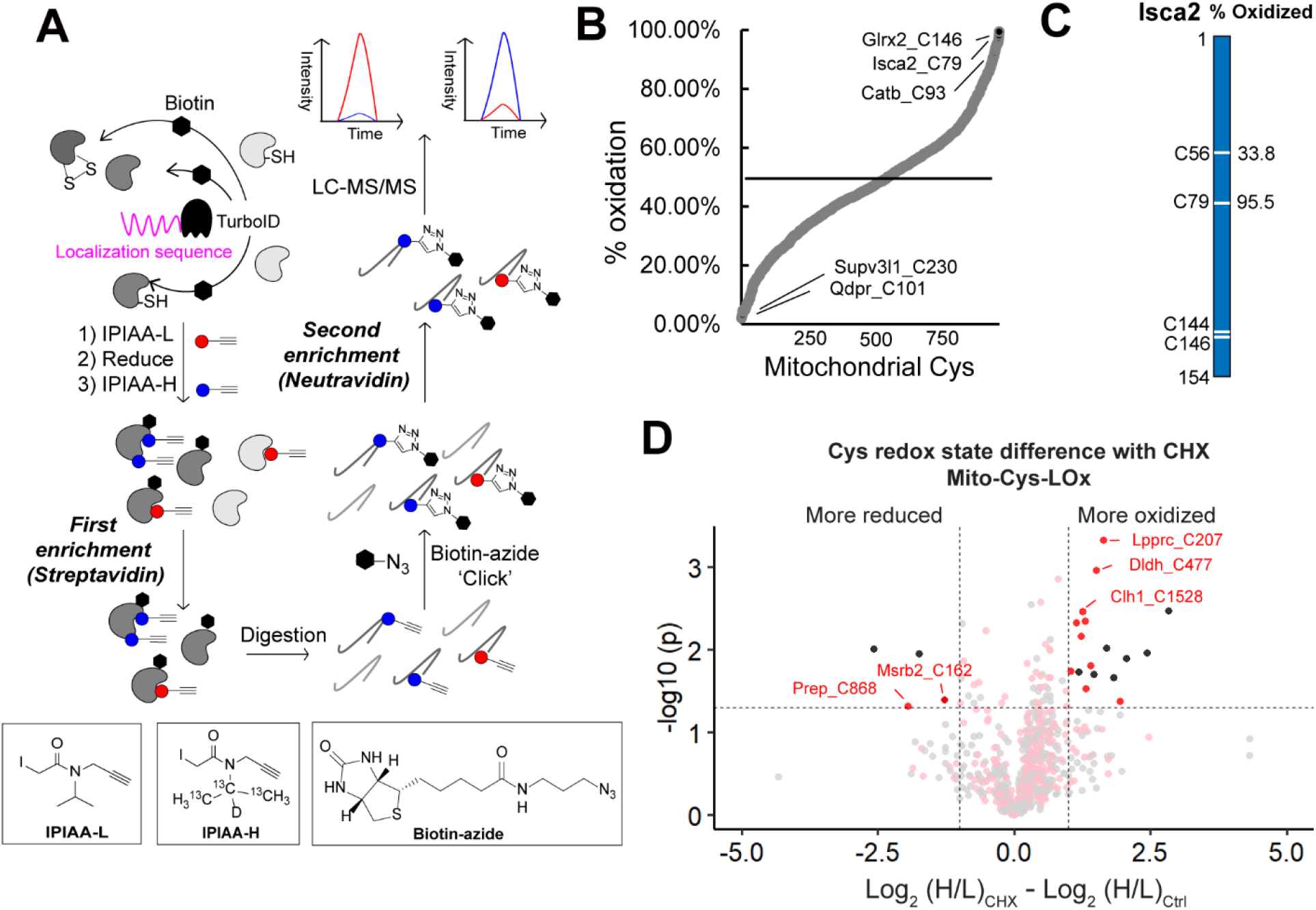
Establishing the Local Cysteine Oxidation (Cys-LOx) method to analyze basal mitochondrial cysteine oxidation states. **A)** Scheme of Cys-LOx method. **B)** Percent oxidation state of mitochondrial cysteines identified with mito-Cys-LOx. **C)** Percent oxidation state of cysteines quantified in an exemplary mitochondrial protein. **D)** Difference in redox states of cysteines quantified with Mito-Cys-LOx with or without dialyzed FBS and CHX treatment. Red dots indicate cysteines localized in mitochondria. Black dots indicate cysteins localized in organelles other than mitochondria. Dialyzed FBS (Dia-FBS) treatment was 36 h and CHX treatment was 100 ug/mL for 6 h at 37 °C. Experiments were performed in triplicate in iBMDM cells. All MS data can be found in **Table S4**.

Cys-LOx analysis of mito-TurboID expressing HEK293T cells quantified 888 total mitochondrial cysteines out of 2739 total cysteines. Of these, 182 had elevated ratios (Log_2_(H/L) > 1), consistent with % oxidation > 50% (**Figure S10A**). Exemplary oxidized cysteines included C129 of membrane-associated progesterone receptor component 1 PGRMC1, C47 of peroxiredoxin-6 PRDX6, C229 of peroxiredoxin-3 PRDX3, which have all been reported as either disulfide or redox centers^82–84^. Reduced cysteines included C46 of Parkinson disease protein 7 PARK7, which was reported as redox sensitive^85^. The median cysteine oxidation state was found to be 34.6%. In the exemplary mitochondrial protein aldehyde dehydrogenase 1 family member B1 ALDH1B1, which plays a major role in the detoxification of alcohol-derived acetaldehyde^86^, the oxidation states of 3 cysteines have been quantified (**Figure S10B**). Both C179 and C386 were found to be > 50% oxidized. By contrast, C169 was calculated to be 9.5% oxidized, indicatively of highly reduced state.

Given our aforementioned demonstration of CHX-enhanced Cys-Loc mitochondrial cysteine specificity, we next incorporated the CHX treatment into the Cys-LOx workflow. Given that CHX treatment can cause cellular stress^87,88^, we first assessed whether translational arrest would impact cysteine oxidation state. By comparing Cys-LOx samples with and without CHX, we found that out of 935 total cysteines quantified, only 5 mitochondrial cysteines were sensitive to CHX treatment (becoming either more oxidized (3) or more reduced (2)) (**Figure S10C**). The reduced cysteines are C419 of mitochondrial-processing peptidase subunit beta PMPCB and C76 of coiled-coil-helix-coiled-coil-helix domain-containing protein 1 CHCHD1 and more oxidized cysteines C218 of pyruvate dehydrogenase Ei component subunit alpha PDHA1, C714 of calcium independent phospholipase A2-gamma PNPLA8 and C428 of mitochondrial-processing peptidase subunit beta PMPCB. Notably PNPLA8 and PMPCB are annotated as having multiple subcellular locations. While we cannot easily rule out direct CHX-induced cysteine oxidation, likely alternative explanations for the observed CHX-dependent changes in cysteine oxidation are subcellular localization-dependent differences in cysteine oxidation together with accumulation of oxidized cysteines in more long-lived proteins.

While we, gratifyingly, observed limited CHX-induced cysteine oxidation, we opted to still proceed conservatively with CHX incorporation into Cys-LOx, given the potential for phenotypic changes and cytotoxicity in response to translation arrest. Within the context of mitochondrial physiology, while CHX treatment is not known to inhibit mitochondrial translation^89^, it does afford decreased gluconeogenesis^87^. To facilitate identification of high confidence mitochondrial oxidation state measurements, we established a “Safe List,” which is comprised of cysteines insensitive to CHX, supporting high confidence mitochondrial-localization not impacted by newly translated mito-TurboID activity. Our Safe List featured 456 total mitochondrial cysteines (**Figure 3I**, gray and pink dots; **Table S4**).

Given our interest in mitochondrial cysteines sensitive to LPS-induced inflammation, we next extended the Cys-LOx method to assay basal redox states of cysteines within mitochondria of immortalized bone marrow derived macrophages (iBMDMs). We opted to use murine, rather than human BMDMs given the established body of literature demonstrating substantial iNOS induction and ROS production by mouse macrophages relative to human macrophages subjected to the same stimuli^90,91^. iBMDMs Cys-LOx analysis of iBMDMs quantified 1,156 total mitochondrial cysteines out of 2,998 total cysteines, in the absence of any stimuli. Of these, 182 had elevated ratios (>1) and % oxidation > 50%, indicative of oxidation (**Figure 4B**). Exemplary oxidized cysteines included C93 of cathepsin B CATB, which has been annotated as a disulfide, C79 of Iron-sulfur cluster assembly 2 Isca2, which is reported as involved in iron-sulfur clusters^92^ and C146 of Glutaredoxin-2 Glrx2, a glutathione-dependent oxidoreductase that facilitates the maintenance of mitochondrial redox homeostasis^93^. Most reduced cysteines included C230 of ATP-dependent RNA helicase Supv3l1 and C101 of dihydropteridine reductase Qdpr, with a percentage oxidation as low as 2.2%. Iron-sulfur cluster assembly 2 homolog Isca2 is an exemplary mitochondrial protein with 2 out of 4 cysteines quantified. Isca2 is involved in the iron-sulfur cluster assembly pathway. Notably, C79 in Isca, which is known to be sensitive to oxidative stress and important for reactivating aconitase^94^, was 95.5% oxidized (**Figure 4C**).

As with our Cys-LOx analysis of HEK239T cells, integration of CHX into the Cys-LOx analysis of iBMDMs revealed minimal CHX-induced changes to measured oxidation states of mitochondrial cysteines. Of the 852 total cysteines quantified across samples, only 12 mitochondrial cysteines were identified as CHX-sensitive, with ten residues showing increased ratios and two decreased ratios, indicative of increased vs decreased oxidation, respectively (**Figure 4D**). As with the HEK293T cysteine redoxome, we also generated a Safe List for the iBMDM mitochondrial cysteine redoxome, which featured 463 total cysteines not impacted by newly translated TurboID (**Table S4**).

### Cys-LOx outperforms SP3-Rox for quantification of LPS-induced changes to cysteine oxidation in immortalized bone marrow derived macrophages (iBMDMs)

Having established the Cys-LOx method, we set out to apply our technology to identify macrophage mitochondrial cysteines sensitive to lipopolysaccharide (LPS) and interferon-gamma (IFNγ)-induced activation. We opted to simultaneously stimulate with LPS+IFNγ given the established synergistic effects both stimuli have on NO production^95–97^. iBMDMs were either vehicle-treated or activated with LPS+IFNγ for 24 hours. As expected, LPS+IFNγ treatment ablates mitochondrial respiration (**Figure 5A, S11**). Additionally, expression of both inducible nitric oxide synthase and other pro-inflammatory genes increased upon treatment, with a more substantial increase observed for cells treated with both cytokines (**Figure 5B, S12**).

**Figure 5.**
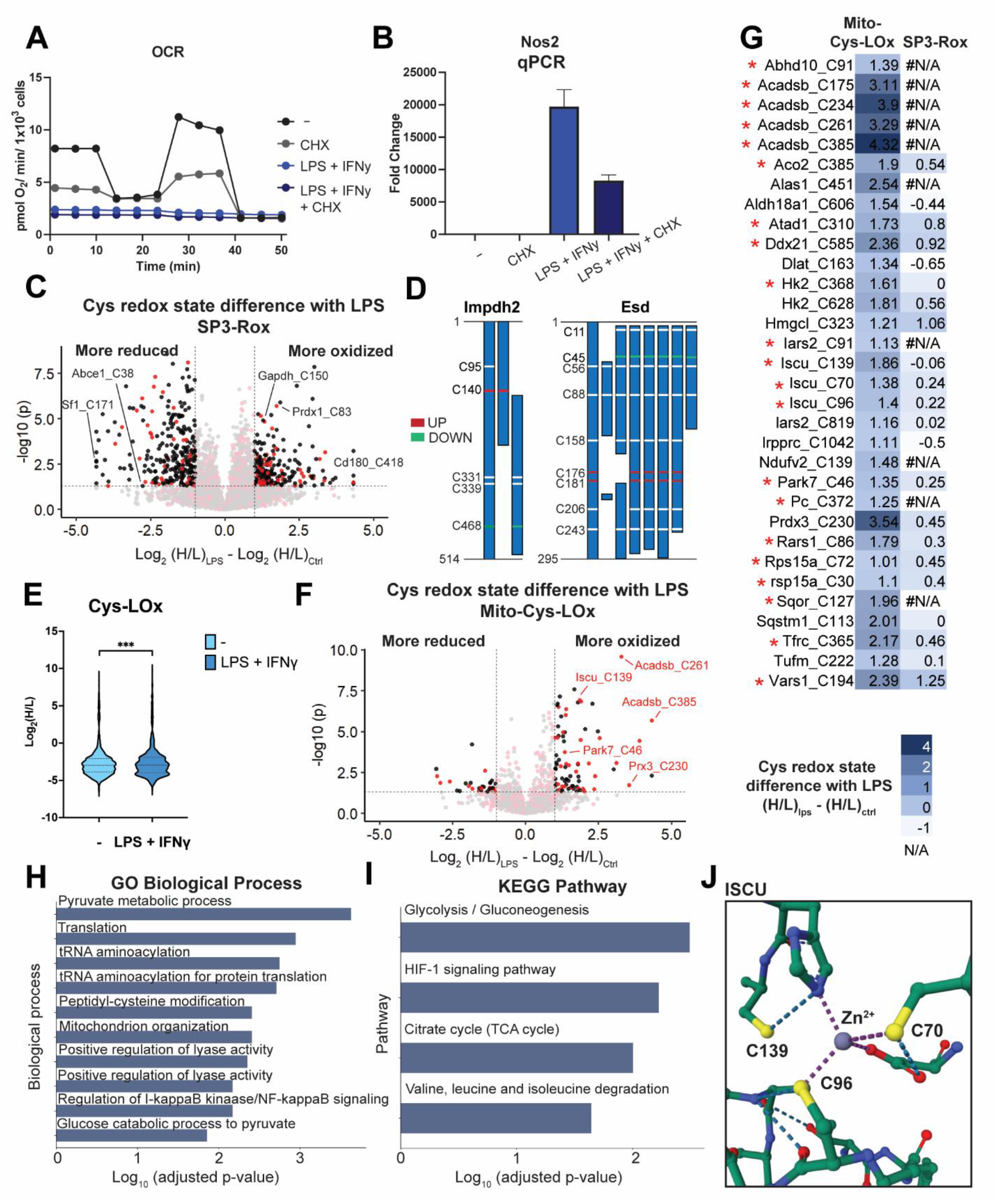
Cys-LOx outperforms SP3-Rox for quantification of LPS-induced changes of mitochondrial cysteine oxidation states. **A)** Oxygen consumption rate (OCR) of mitochondria for control, LPS+IFNγ, CHX or both. This is representative trace of one biological replicate with 5 technical replicates. For some timepoints, the symbols obscure the error bars. **B)** qCPR analysis of expression of Nos2 with control, LPS+IFNγ, CHX or both. **C)** Difference of redox states for cysteines quantified with SP3-Rox with or without LPS+IFNγ treatment. Red dots indicate cysteines localized in mitochondria. Black dots indicate cysteins localized in organelles other than mitochondria. **D)** Different redox states of cysteines quantified with SP3-Rox with or without LPS+IFNγ treatment in exemplary proteins with different splice forms. **E)** Redox states of cysteines quantified with Mito-Cys-LOx with or without LPS+IFNγ treatment. Statistical significance was calculated with paired Student’s t-tests, **** p<0.001. **F)** Difference of redox states for cysteines quantified with Mito-Cys-LOx with or without LPS+IFNγ treatment. Red dots indicate cysteines localized in mitochondria. Black dots indicate cysteins localized in organelles other than mitochondria. **G)** Difference of redox states for cysteines quantified with Mito-Cys-LOx and SP3-Rox with or without LPS+IFNγ treatment. * indicates cysteines in the Safe List we generated that are insensitive to translational arrest. **H)** GO biological process analysis of mitochondrial cysteines quantified with mito-Cys-ROx that showed more oxidized redox states upon LPS+IFNγ treatment. **I)** KEGG pathway analysis of mitochondrial cysteines quantified with mito-Cys-ROx that showed more oxidized redox states upon LPS+IFNγ treatment. **J)** Crystal structure of ISCU with cysteines more oxidized with LPS+IFNγ treatment (PDB 1WFZ). Experiments were performed in triplicates and technical replicates in iBMDM cells. Dialyzed FBS (Dia-FBS) treatment was 36 hrs, CHX treatment was 100 ug/mL for 6 h at 37 °C and LPS+IFNγ treatment was 100 ng/mL LPS and 20 ng/mL IFNγ for 24 h at 37 °C. All MS data can be found in **Table S5**.

Having previously established the SP3-Rox method^11^, which reports proteome-wide cell-state-dependent changes to cysteine oxidation, we next subjected the iBMDMs to SP3-Rox analysis. Our goals were to establish a baseline for LPS-induced whole-cell cysteine oxidation from which we could compare the Cys-LOx method, and to assess whether the Cys-LOx method could capture mitochondrial cysteines not readily quantified by established, bulk proteomic methods. Additionally, we were optimistic that Cys-LOx would reveal mitochondrial specific redox changes that get masked by the bulk analysis of SP3-Rox. In aggregate, SP3-Rox analysis captured 7,523 total cysteines and identified 290 cysteines that showed increased oxidation upon LPS+IFNγ treatment (**Figure 5C**). Quantified cysteine ratios were significantly increased after LPS+IFNγ treatment, indicative of substantial cell-wide oxidation (**Figure S13**). This finding is to be expected given the marked increase in NOS expression and resulting widespread cysteine nitrosylation. Overall, 1742 total mitochondrial cysteines were identified and 70 were found to exhibit increased oxidation in response to LPS+IFNγ (**Figure 5C**). Of the LPS+IFNγ-oxidation-sensitive cysteines identified, several have been previously identified as redox active, including C150 of glyceraldehyde-3-phosphate dehydrogenase Gapdh, C147 of superoxide dismutase [Cu-Zn] Sod1, C54 of peroxiredoxin-4 Prdx4, C83 of peroxiredoxin-1 Prdx1, and C1557 of fatty acid synthase Fasn^11,24^.

Curiously, we also observed a marked population of 293 total cysteines with significantly reduced oxidation state upon LPS+IFNγ treatment, as analyzed by SP3-Rox of bulk proteomes. Given the large burst of ROS and RNS produced upon macrophage stimulation, this cysteine population was unexpectedly large. Consistent with prior reports^98,99^, we see a decrease in total glutathione concentration upon activation with LPS + IFNγ, excluding a reducing environment from rationalizing the reduced subset (**Figure S14A**). Pathway analysis for the reduced population revealed a marked enrichment for genes involved in RNA and protein biogenesis, including splicing, NFkB gene targets, and translation in both Gene Ontology (GO) and Kyoto Encyclopedia of Genes and Genomes (KEGG) pathway analysis (**Figure S14B, S14C**). We postulated that post-stimulation, translation of LPS+IFNγ responsive genes was likely responsible for the observed population of highly reduced cysteines. Supporting this finding, we observed a comparatively short (2h) half-life protein ABCE1 (**Figure S14D**) that contained more reduced cysteines C38 and C65, with oxidation states reduced from 80.7% to 39.5% and 87.8% to 58.7%, respectively upon LPS+IFNγ treatment. Further implicating altered gene expression, we observed 14 proteins which harbored cysteines with both increased and decreased oxidation states, including Inosine-5’-monophosphate dehydrogenase 2 (Impdh2 C140, C468), S-formylglutathione hydrolase (Esd C176, C181, C45), and Valine–tRNA ligase (Vars C194, C1051) (**Figure 5D, S14E**), all of which encode multiple splice forms, suggestive that LPS+IFNγ-dependent alternative splicing may play a role in production of this population of proteins (**Table S5**). Gratifyingly, only 49 mitochondrial cysteines were found to be more reduced, indicating the mitochondrial proteome is relatively insensitive to the observed global reducing response to LPS+IFNγ.

Application of Mito-Cys-LOx to our LPS+IFNγ iBMDM system quantified, in aggregate, 1,451 total and 559 mitochondrial cysteines. Similar to SP3-Rox, a marked increase in the ratios of identified cysteines was observed after LPS+IFNγ treatment (**Figure 5E**). 32 mitochondrial cysteines exhibited significant increase in the H:L ratio upon LPS treatment (Log_2_(H/L)_LPS_ - Log_2_(H/L)_Ctrl_ > 1), indicating increased oxidation (**Figure 5F, 5G**). 23 of the 32 cysteines that showed increased oxidation state in the Cys-LOx analysis were identified in the SP3-ROx analysis. Of these, only 2 residues (Hmgcl_C323 and Vars1_C194) also showed increased oxidation upon LPS+IFNγ activation in the SP3-ROx analysis, while the oxidation state of the other 21 was not significantly changed (**Figure 5G**). This result highlights the value of subcellular redox state analysis in capturing compartment-specific changes in cysteine oxidation, as a global analysis masks the redox changes that are specific to a single compartment.

We observed a pronounced decrease in mitochondrial respiration following 6h CHX treatment, substantiating our prior concerns for potential CHX-dependent alterations in mitochondrial cysteine redox states. (**Figure 5A, S7**). Additionally, qPCR analysis of genes associated with response to LPS+IFNγ revealed both comparatively CHX-insensitive (*Irg1* and *Il1b*) and highly CHX-sensitive (Tfna and Il6) changes to gene expression (**Figure S8**). We referenced our previously generated iBMDM cysteine safe list (**Table S4**) to delineate high confidence mitochondrial cysteines that exhibit increased oxidation in response to LPS+IFNγ. Within the 32 mitochondrial cysteines exhibiting more elevated H:L ratios, indicating increased oxidation upon LPS treatment, we identified 21 of them to be insensitive to translational arrest, which are indicated by the red asterisks (**Figure 5G**).

Pathway analysis of the oxidized subset revealed peptidyl cysteine modification (GO: 0018198) as a major enriched Gene Ontology biological process (**Figure 5H**), with glycolysis (while a cytosolic process, this enrichment stems from oxidation of cysteines in hexokinase, which is tethered to the outer mitochondrial membrane^100^) and HIF-1 signaling as significantly enriched KEGG pathways (**Figure 5I**). Supporting the robustness of our LPS+ IFNγ dataset, many of the cysteines identified by Cys-Lox are residues previously characterized as sensitive to oxidative modification. Examples include Iron-sulfur cluster assembly enzyme Iscu C139 and C70 (cysteine persulfide)101, Parkinson disease protein 7 homolog Park7 C46 (cysteine palmitoyl)^85^, Hydroxymethylglutaryl-CoA lyase Hmgcl C323 (disulfide)^102^, Thioredoxin-dependent peroxide reductase Prdx3 C230 (disulfide)^103^, and Aconitate hydratase Aco2 C385^48^. All three cysteines we identified as sensitive to LPS+IFNγ in Iscu are proximal to or serve as ligands for the coordinated Zn^2+^ ion (**Figure 5J**), suggesting that these residues may play a role in sensing oxidative stress. Intriguingly, C385 of Aco2 binds the active site Fe-S cluster that is essential for catalytic activity, and oxidation of this cysteine has been suggested to render the protein inactive^48,104^. The result suggested that modification of aconitase – in addition to those of isocitrate dehydrogenase and succinate dehydrogenase^105^ – may result in cellular respiration defects caused by pro-inflammatory activation. We additionally identified a number of LPS+IFNγ sensitive cysteines that have not been previously annotated as sites of oxidative modification (e.g. C96 in Iron-sulfur cluster assembly enzyme Iscu and C91 and C819 in Isoleucine--tRNA ligase Iars2), which likely represent novel sites of redox regulation. Taken together these residues, highlights the the general utility of Cys-Lox for studying the subcellular cysteinome, including in the context of mitochondrial sensitivity to cellular inflammatory processes.

## Discussion

Here we establish two novel cysteine chemoproteomic platforms, Local Cysteine Capture (Cys-LoC) and Local Cysteine Oxidation (Cys-LOx), which enable compartment-specific capture of cysteines and quantification of changes to local cysteine oxidation state, respectively. Both Cys-LoC and Cys-LOx implement a customized two-step biotinylation workflow that features sequential enrichment of subcellular-localized proteins (e.g. cytosol, ER, golgi, mitochondria, and nucleus, as labeled by targeted TurboID constructs) followed by peptide-level click-conjugation biotinylation and a subsequent second round of enrichment to capture cysteine-containing peptides derived from the TurboID-labeled proteins. While our method is in many ways conceptually similar to TurboID-based subcellular phosphoproteomics^59^, to our knowledge, such sequential rounds of biotin-avidin capture, first at the protein level and then at the peptide level, have not been reported previously. Application of Cys-LOx to iBMDMs quantified 559 total mitochondrial cysteines and 32 sensitive to LPS+IFNγ treatment. Notably, a number of these residues are found in proteins involved in iron sulfur cluster biogenesis and respiration, including aconitase and iron-sulfur cluster assembly enzyme ISCU. Given the importance of these proteins to mitochondrial function, we expect that oxidative modifications to some of these cysteines may, in part, explain the marked respiration defects caused by LPS+IFNγ (**Figure 5A**).

Showcasing the utility of Cys-LOx, we found that bulk SP3-Rox proteome cysteine redox analysis of LPS+IFNγ stimulated iBMDMs was complicated by the appearance of a substantial fraction of cysteines that showed decreased ratios, indicative of decreased oxidation, after treatment. Our KEGG/GO analyses, together with identification of cysteines likely impacted by alternative splicing, points towards LPS+IFNγ-induced transcription as a likely cause of this curious appearance of more reduced residues after oxidative assault. As the SP3-Rox method relies on a ratio difference calculation, it is relatively insensitive to protein abundance changes. However, in this case, we expect that turnover of NO-damaged protein together with production of newly synthesized protein could in part rationalize the ratios observed. This model is consistent with prior studies that reported substantial remodeling of the macrophage proteome in response to IFNγ+LPS^106^.

As mitochondrial isolation is feasible using established biochemical fractionation methods, we expect that some of our findings could be corroborated by fractionation-based OxiCat platforms^34,35^. Looking beyond organelles that can be separated by density gradient centrifugation methods, we expect that the Cys-LoC and Cys-LOx methods will prove most useful for chemoproteomic analysis of subcellular compartments, protein complexes, and cell types that are not readily amenable to biochemical fractionation. Examples of such applications include membraneless organelles, such as stress granules and P-bodies^41,42,107^, and proteins localized to neuronal axons^108^; both of which have successfully implemented proximity labeling methods to build their proteomes Fully realizing the utility of Cys-LoC and Cys-LOx in such applications may require methodological optimization to further increase cysteine coverage, which will be of particular value for redox analyses that require high cysteine coverage to calculate difference values between treated and control groups.

One challenge that we encountered while establishing the Cys-LoC and Cys-LOx methods was the seemingly low compartment specificity of proteins and cysteines captured by TurboID proximity labeling. Through our in-depth analysis of sources of aberrant TurboID performance, we first pinpointed endogenous biotin derived from serum as a source of non-compartment specific labeling, consistent with prior studies^69,72^. While depletion of endogenous biotin using dialyzed serum afforded a modest increase in subcellular specificity, a more pronounced effect was observed after treatment with cycloheximide. These findings are consistent with newly translated TurboID as a major source of the observed seemingly non-specific biotinylation. CHX treatment proved particularly useful in our study for establishing Safe Lists of cysteines not impacted by this unwanted TurboID activity. As shown by the toxicity of CHX to mitochondrial respiration, both measured in our study and reported previously^87^, CHX likely will prove suboptimal as a universal solution to the TurboID localization conundrum. Further exemplifying the need for additional methods, we also found that the CHX treatment afforded the greatest improvement in localization specificity for the mito-TurboID construct, with more modest enhancement observed for other compartments analyzed. While a clear rationale for these construct-dependent differences remains to be fully explored, we expect that the previously reported CHX-dependent impact on subcellular RNA localization, as assayed by APEX-Seq^109^, may implicate a confluence of RNA- and protein-localization factors.

Looking beyond CHX, we can envision that two-step cysteine capture methods with enhanced performance could be achieved using several relevant alternative approaches, including, for example, the previously reported Split TurboID^40^, TurboID caged using unnatural amino acid (UAA) incorporation^110^ and potentially alternative genetic approaches that afford tighter control over protein expression (e.g. Flp-In™ T-Rex™ system; Invitrogen Life Technologies) that would reduce the fraction of newly synthesized protein. As BioID is less active^38^, we expect that the newly translated BioID proteins should less substantially impact proximity labeling performance. However, this low catalytic activity may prove insufficient to achieve high coverage two-step cysteine capture. Furthermore, as shown by our absolute quantitation of mitochondrial and whole cell biotin concentrations achieved after exogenous biotin addition, limited biotin entry into organelles may continue to confound such improved platforms. Alternative proximity labeling methods that utilize reagents with enhanced membrane solubility may be required.

## Methods

### Cloning of different TurboID constructs

List of plasmids with detailed information used in this study can be found in **Table S6**.

### Biotinylation with TurboID

For transiently expressed TurboID, biotin labeling was initiated 24 h after transfection. Biotin was directly added to cells at a final concentration of 500 μM and incubated for 1 h at 37 °C. After washing with cold DPBS for 3 times, cells were harvested by centrifugation (4,500 *g*, 5 min, 4 °C), washed twice with cold DPBS, lysed in RIPA buffer, and clarified by centrifuging (21,000 *g*, 10 min, 4 °C).

### Proteomic sample preparation for TurboID, Cys-LoC and Cys-LOx

Biotinylated lysates (500 μL of 1 mg/mL, prepared as described above) were labeled with either 2 mM IAA for Cys-LoC or 2 mM IPIAA-L in 2M urea for Cys-LOx for 1h at RT. 50 uL pierce streptavidin agarose beads were washed with RIPA and incubated with lysates for 2 h at RT. The proteins bound to beads were washed, reduced with 1 mM DTT for 15 min at 65 °C, and labeled with 2 mM IAA for Cys-LoC or 2 mM IPIAA-H for Cys-LOx for 1h at RT. Then, proteins were digested off the bead overnight at 37 °C with trypsin. For one-step TurboID workflow, peptides were desalted with C18 column and analyzed by LC-MS/MS. For Cys-LoC and Cys-LOx, after digestion, CuAAC was performed with biotin-azide for 1h at RT. SP3 sample clean up was performed as reported^11,25,111^. Peptides were eluted from SP3 beads with 200 μL of 2% DMSO in water for 30 min at 37 °C. For each sample, 50 μL of NeutrAvidin Agarose resins were washed and incubated with the eluted samples for 2 h at RT. After incubation, the beads were pelleted by centrifugation (21,000 *g*, 1 min) and washed. Bound peptides were eluted twice with 60 μL of 80% acetonitrile in MB water containing 0.1% FA. The first 10 min incubation at RT and the second one at 72 °C. The combined eluants were dried (SpeedVac), then reconstituted with 5% acetonitrile and 1% FA in MB water and analyzed by LC-MS/MS.

### Proteomic sample preparation for SP3-Rox of iBMDMs

iBMDM cells were treated either with or without 100 ng/mL LPS and 20 ng/mL IFNγ for 24 h at 37 °C. Cells were then harvested and SP3-Rox procedure was carried out as reported^11^.

### Liquid-chromatography tandem mass-spectrometry (LC-MS/MS) analysis

The samples were analyzed by liquid chromatography tandem mass spectrometry using a Thermo Scientific™ Orbitrap Eclipse™ Tribrid™ mass spectrometer.

### Protein, peptide, and cysteine identification

Raw data collected by LC-MS/MS were searched with MSFragger (v3.3) and FragPipe (v19.0). For Cys-LoC, Cysteine residues were searched with differential modification C+463.2366. For Cys-LOx and SP3-Rox, MS1 labeling quant was enabled with Light set as C+463.2366 and Heavy set as C+467.2529. MS1 intensity ratio of heavy and light labeled cysteine peptides were reported^112^.

Additional experimental details can be found in the **Supporting Information**.

## Acknowledgements

This study was supported by a Packard Fellowship (K.M.B.), National Institutes of Health **DP2 OD030950-01**(K.M.B.), National Institutes of Health R35GM138003 and P30DK063491 (A.S.D.), W.M. Keck Foundation Award 995337 (A.S.D.), TRDRP T31DT1800 (T.Y.), NIGMS System and Integrative Biology 5T32GM008185-33 (L.M.B.), NIGMS UCLA Chemistry Biology Interface T32GM136614 (A.R.J, M.V., and A.B.B). We thank all members of the Divakaruni and Backus labs for helpful suggestions.

## DATA AVAILABILITY

The MS data have been deposited to the ProteomeXchange Consortium (http://proteomecentral.proteomexchange.org) via the PRIDE^113,114^ partner repository with the dataset identifier PXD039626. File and peptide details are listed in Table S9.

## Author Contributions

T.Y., A.S.D, and K.M.B. conceived of the project. T.Y. A.R.J., M.V., A.E.J. A.B.B, A.C.T, and S.L.Y collected data. T.Y, A.R.J., M.V., A.E.J. A.B.B, L.M.B. and H.S.D performed data analysis. L.M.B and H.S.D wrote software. T.Y., A.R.J., M.V., A.E.J. A.B.B, contributed to figures. T.Y. and K.M.B wrote the manuscript with assistance from all authors.

