## Supplementary Information for "Proximity-labeling chemoproteomics defines the subcellular cysteinome and inflammation-responsive mitochondrial redoxome"

### Contents

**(A) Supplementary figures 2**
**(B) Supplementary tables 16**
**(C) Methods 22**
**(D) References 28**

### Supplementary Figures

| Subcellular compartment | Localization sequence |
|---|---|
| Cytosol | LPPLERLTL (C-term) |
| Endoplasmic Reticulum | MDPVVVLGLCLSCLLLLSLWKQSYGGG (N-term) |
| Golgi | FLWRIFCFRK (C-term) |
| Mitochondria | MLATRVFSLVGKRAISTSVCVRAH (N-term) |
| Nucleus | PKKRKV (C-term) |

**Figure S1.** Different localization sequence.

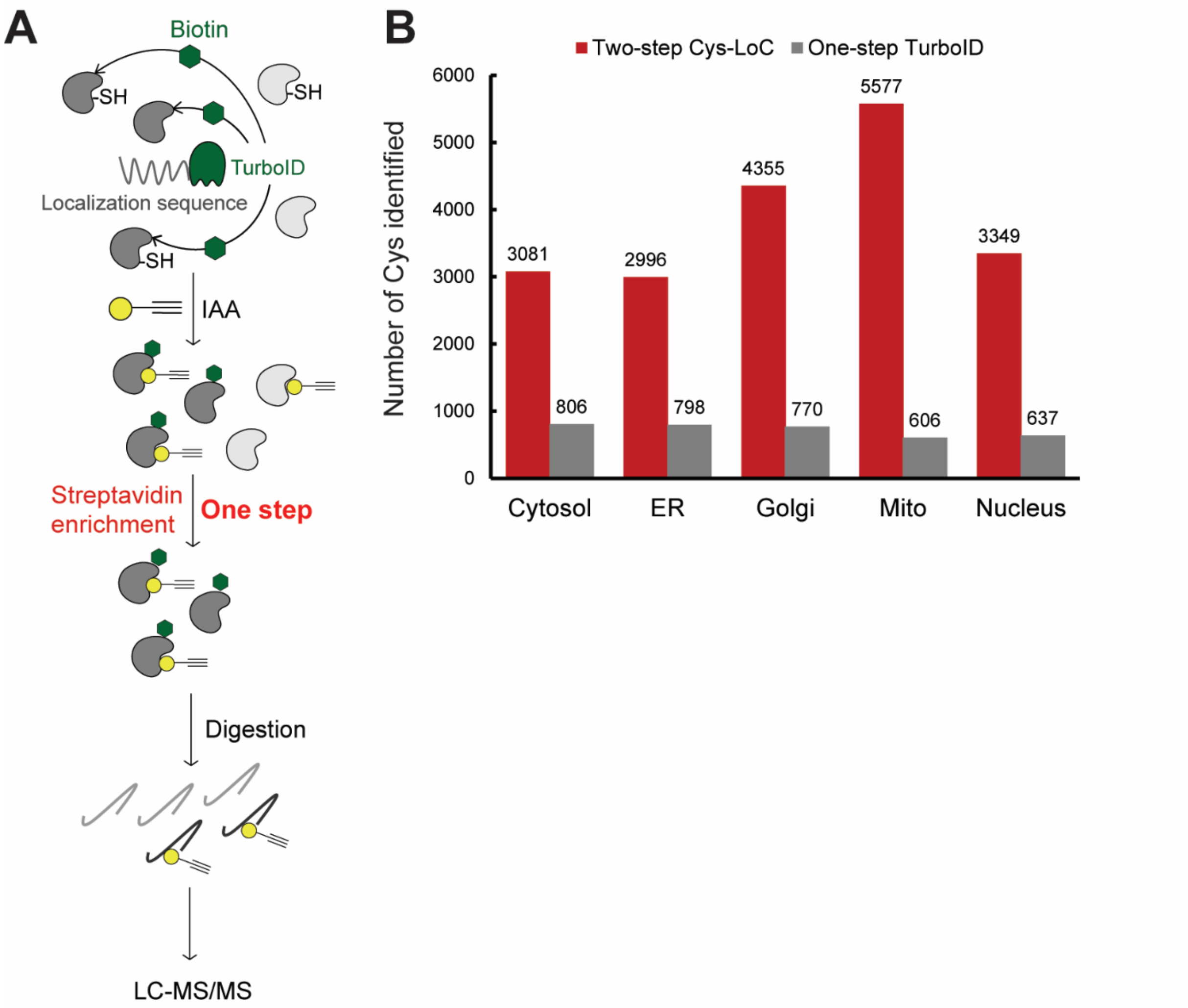


**Figure S2**. **A)** Scheme of One-step TurboID. **B)** Number of cysteines identified by Two-step Cys-LoC vs One-step TurboID in different subcellular compartments. Experiments were performed in singlicate in 293T cells.

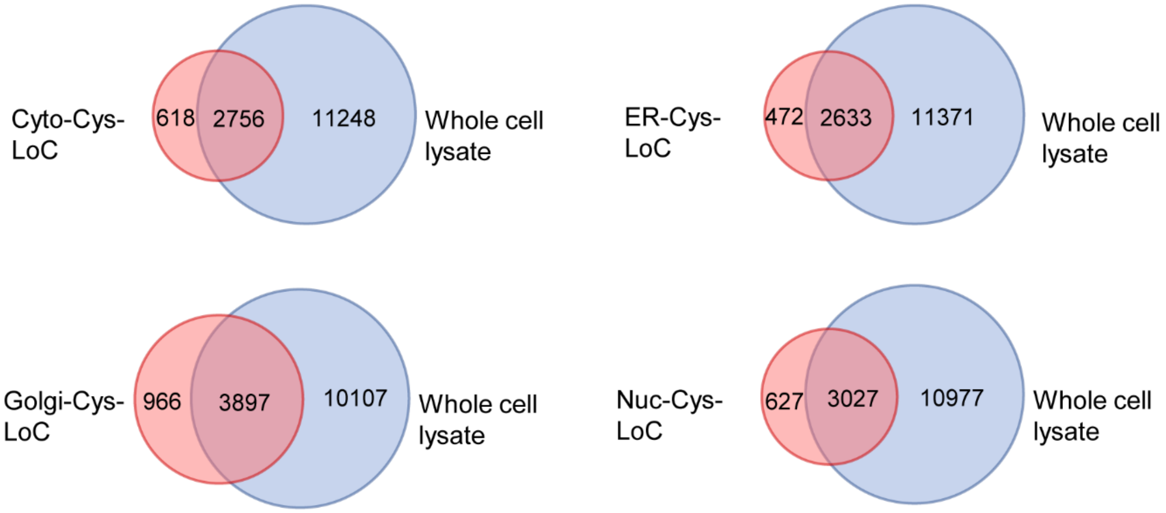


**Figure S3.** Cysteines identified in different subcellular compartments with Cys-LoC. Experiments were performed in duplicate in 293T cells.

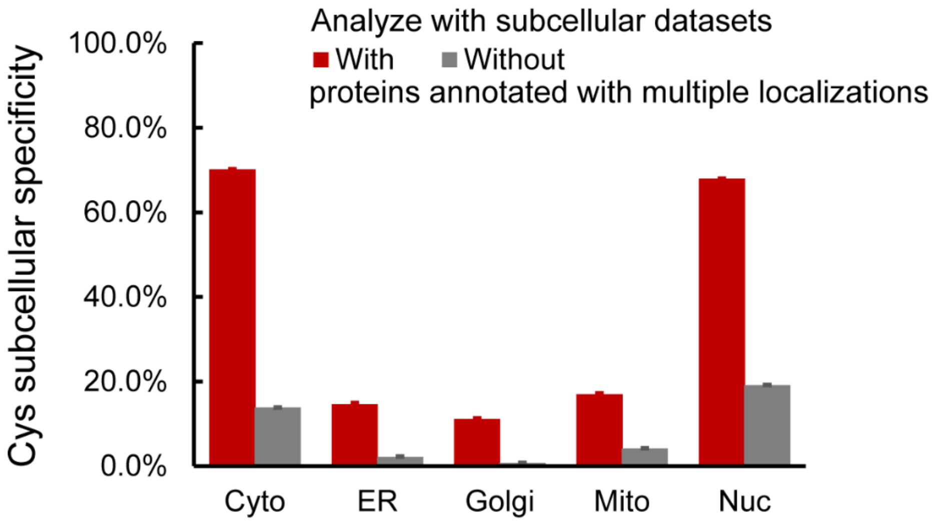


**Figure S4.** Subcellular specificity of proteins identified with TurboID, using localization datasets including or excluding proteins with multiple localizations. Experiments were performed in duplicate in 293T cells.

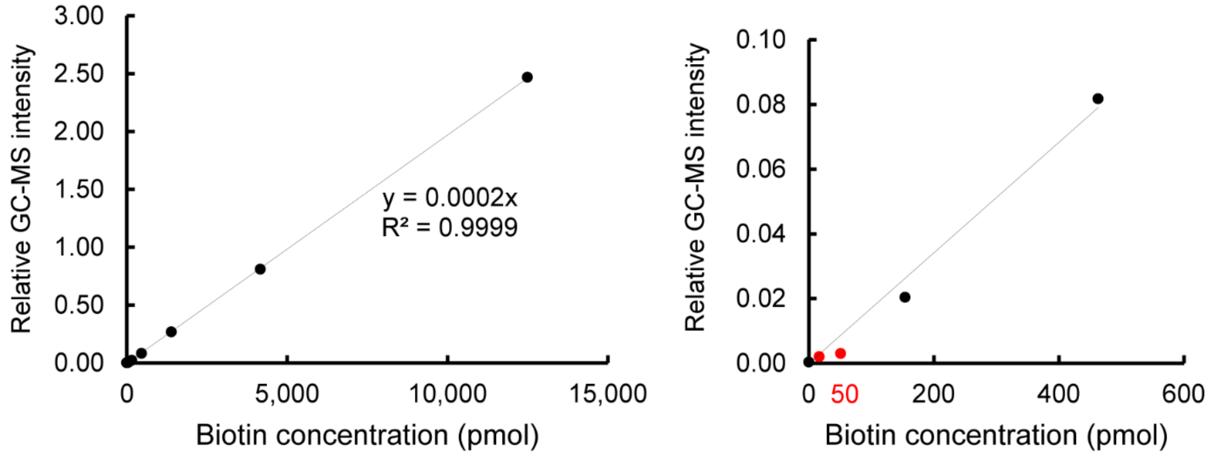


**Figure S5.** Standard curve of biotin molecules detected with optimized GC-MS protocol.

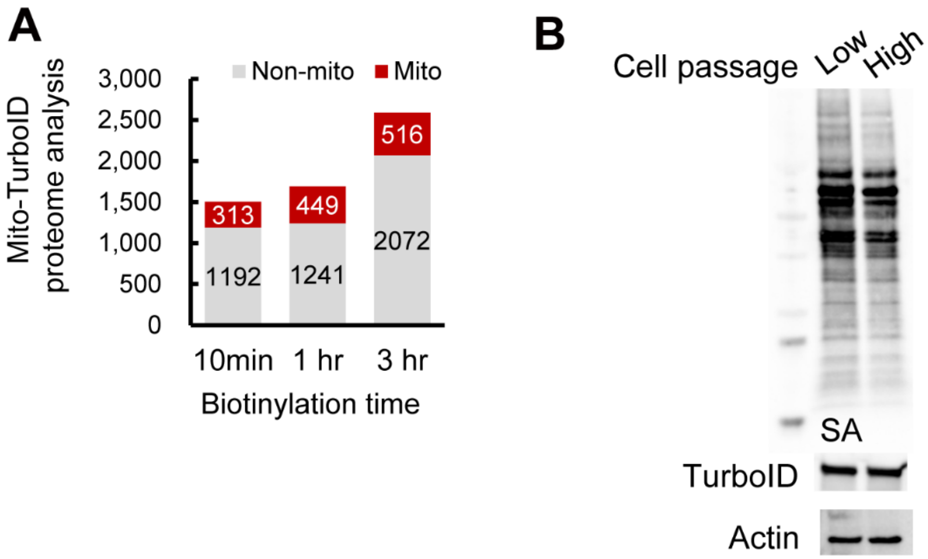


**Figure S6. A)** Number of mitochondrial annotated proteins and non-mitochondrial annotated proteins identified with mito-TurboID with biotinylation of 10 min or 1hr or 3hr. Experiments were performed in singlicate in HEK293T cells. **B)** Biotinylation efficiency of mito-TurboID stably expressed in HEK293T cells < 10 passages (Low) or cells > 10 passages (High).

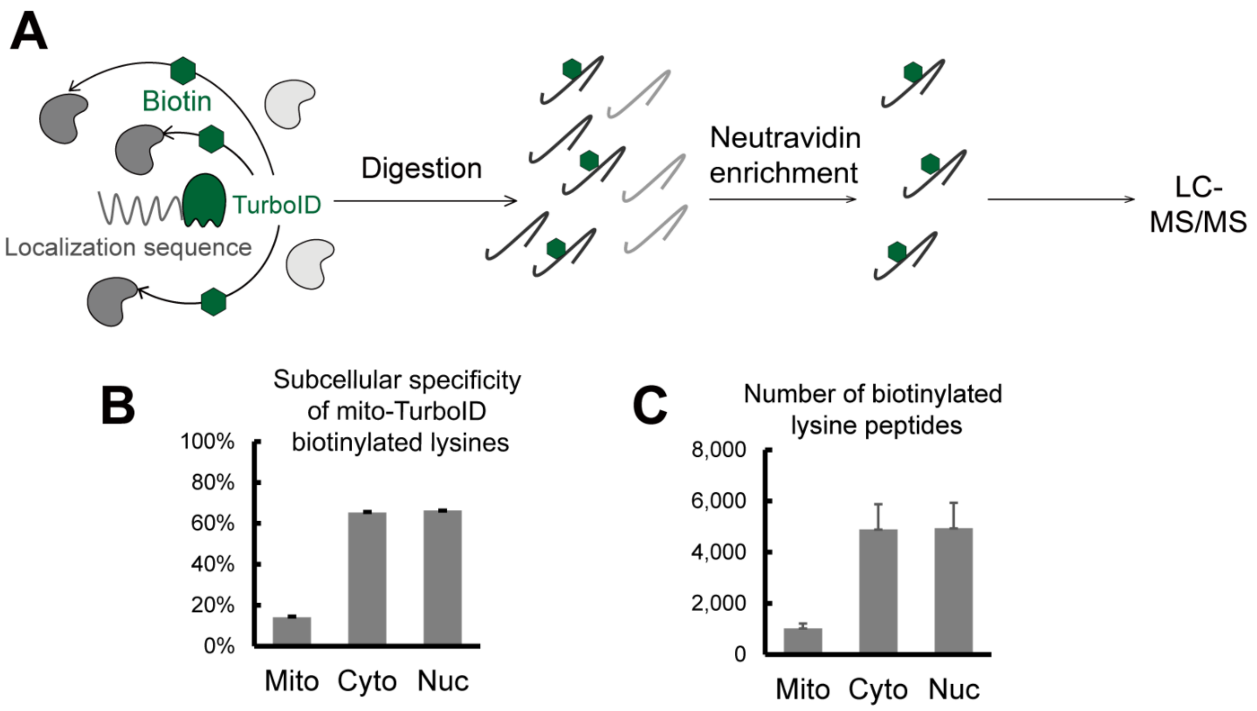


**Figure S7. A)** Scheme of profiling lysines biotinylated by TurboID ligase. **B)** Subcellular specificity of mito-TurboID biotinylated lysines. **C)** Number of biotinylated lysine peptides annotated in different compartments enriched with mito-TurboID. Experiments were performed in duplicate in HEK293T cells.

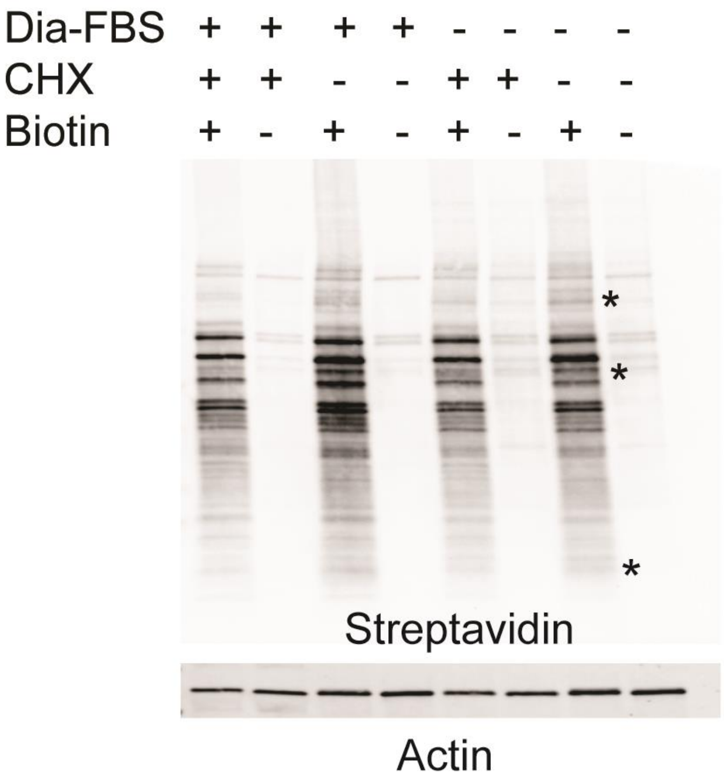


**Figure S8.** Biotinylation of mito-TurboID with no treatment, with dialyzed FBS, with CHX treatment or with both. Dialyzed FBS (Dia-FBS) treatment was 36 h and CHX treatment was 100 ug/mL for 6 h at 37 °C if not specified.

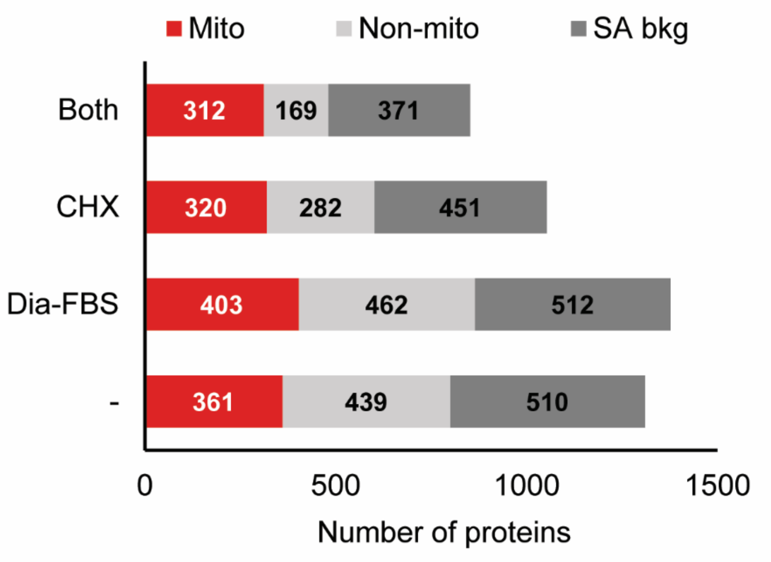


**Figure S9.** Proteome analysis of the proteins enriched with mito-TurboID with control condition, with dialyzed FBS, with CHX treatment or with both. Dialyzed FBS (Dia-FBS) treatment was 36 h and CHX treatment was 100 ug/mL for 6 h at 37 °C if not specified. Experiments were performed in triplicates in HEK293T cells.

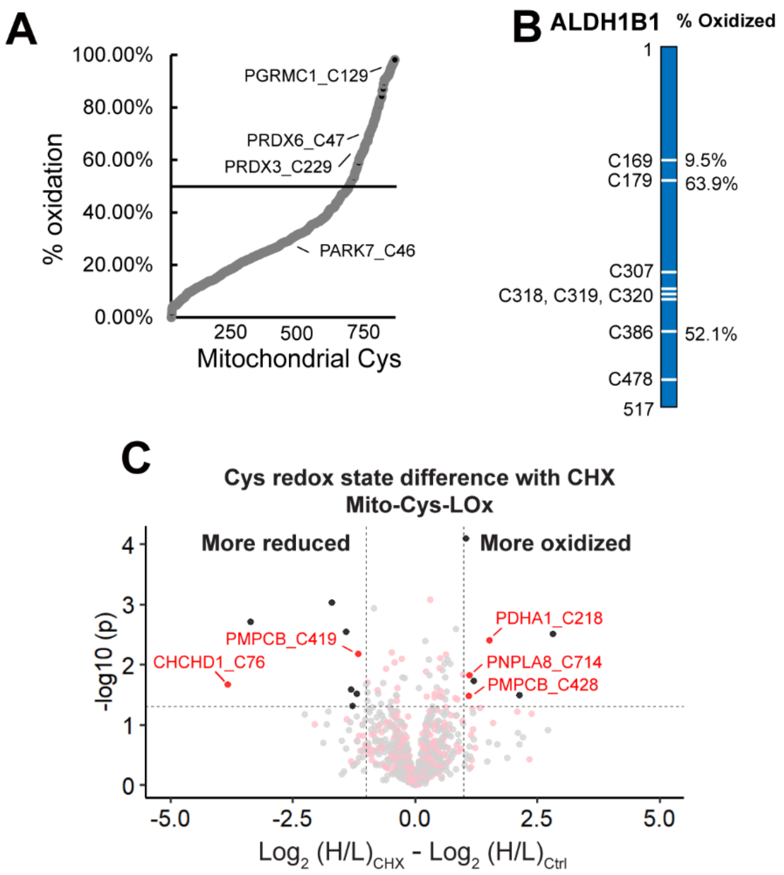


**Figure S10. A)** Percentage oxidation states of mitochondrial cysteines identified with mito-Cys-LOx. **B)** Percentage oxidation states of cysteines quantified in exemplary mitochondrial proteins. **C)** Difference in redox states of cysteines quantified with Mito-Cys-LOx with or without dialyzed FBS and CHX treatment. Red dots indicate cysteines localized in mitochondria. Black dots indicate cysteins localized in organelles other than mitochondria. Dialyzed FBS (Dia-FBS) treatment was 36 h and CHX treatment was 100 ug/mL for 6 h at 37 °C. Experiments were performed in triplicates in HEK293T cells.

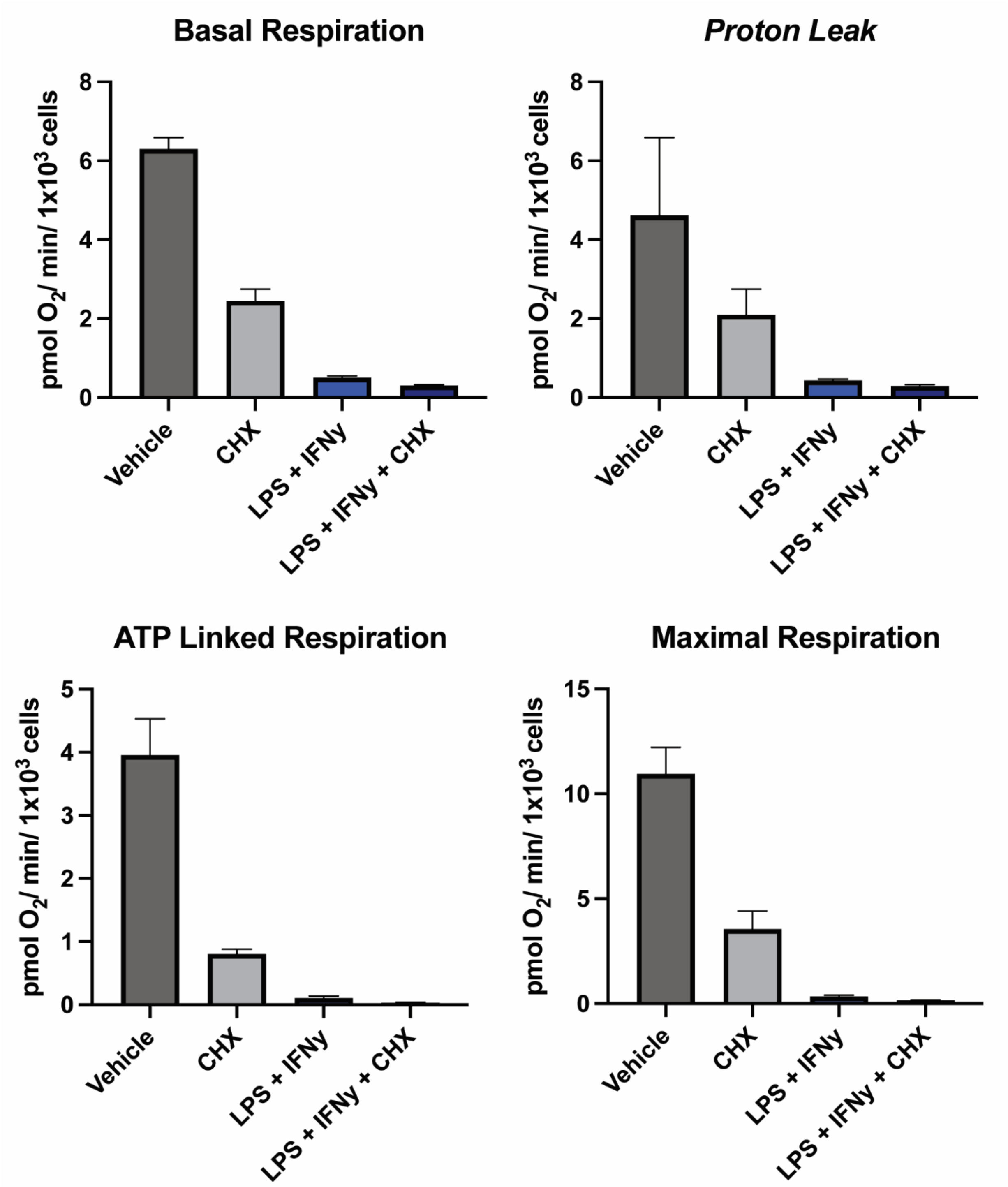


**Figure S11.** Seahorse analysis of iBMDMs with control, LPS+IFNγ, CHX or both. Dialyzed FBS (Dia-FBS) treatment was 36 hrs, CHX treatment was 100 ug/mL for 6 h at 37 °C and LPS+IFNγ treatment was 100 ng/mL LPS and 20 ng/mL IFNγ for 24 h at 37 °C if not specified. Experiments were performed in 5 technical replicates in iBMDM cells.

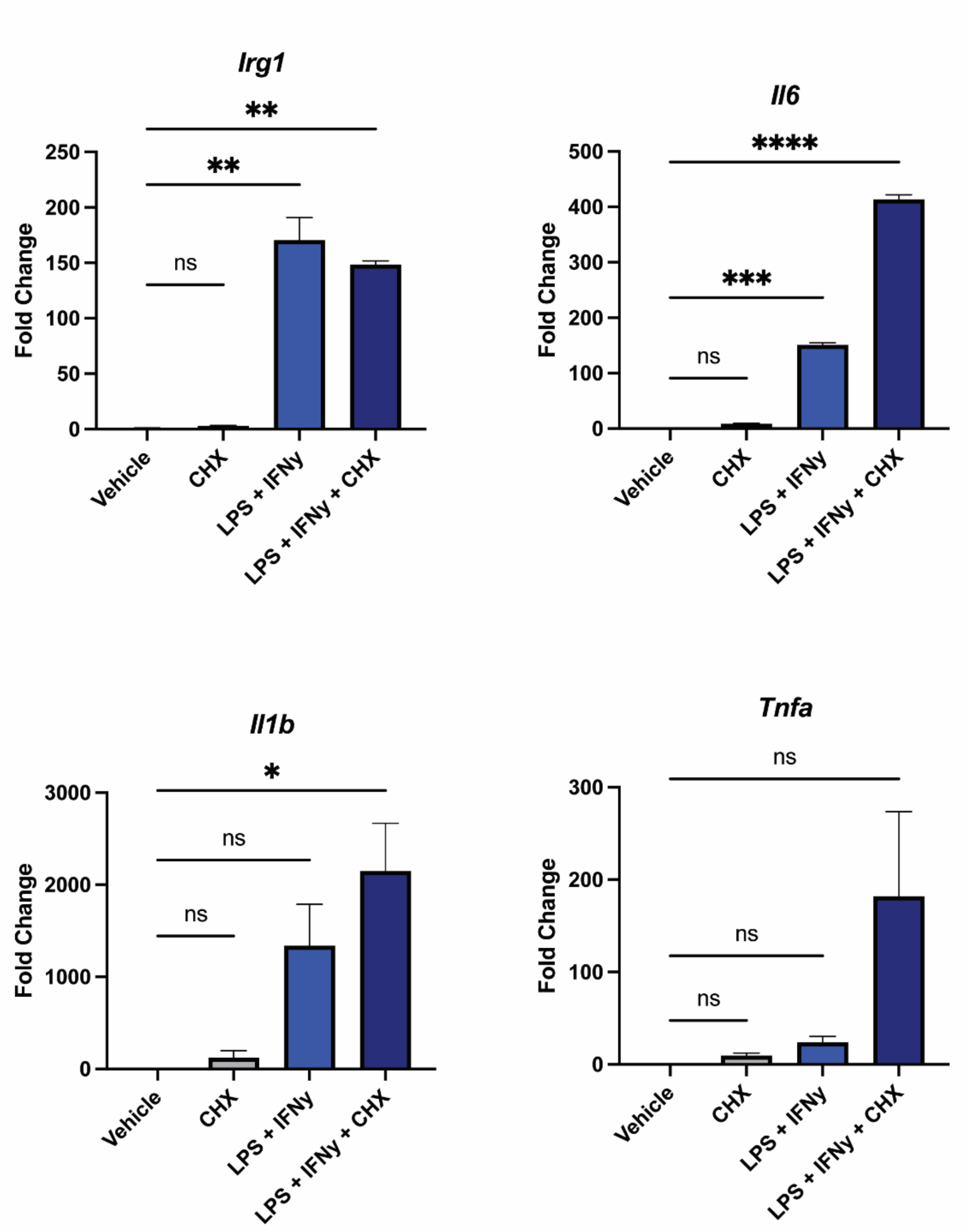


**Figure S12.** qPCR analysis of LPS related genes with control, LPS+IFNγ, CHX or both. Dialyzed FBS (Dia-FBS) treatment was 36 hrs, CHX treatment was 100 ug/mL for 6 h at 37 °C and LPS+IFNγ treatment was 100 ng/mL LPS and 20 ng/mL IFNγ for 24 h at 37 °C if not specified. Statistical significance was calculated with unpaired Student's t-tests, * $p<0.05$, ** $p<0.01$, *** $p<0.005$. NS $p>0.05$. Experiments were performed in triplicates in iBMDM cells.

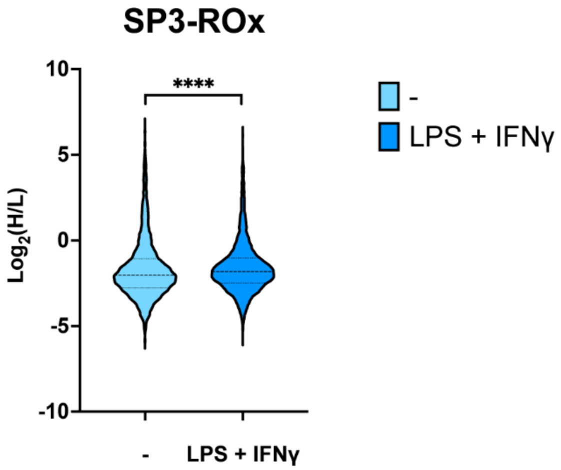


**Figure S13.** Redox states of cysteines quantified with SP3-Rox with or without LPS+IFNγ treatment. LPS+IFNγ treatment was 100 ng/mL LPS and 20 ng/mL IFNγ for 24 h at 37 °C. Experiments were performed in triplicates in iBMDM cells. Statistical significance was calculated with paired Student's t-tests, **** p<0.001.

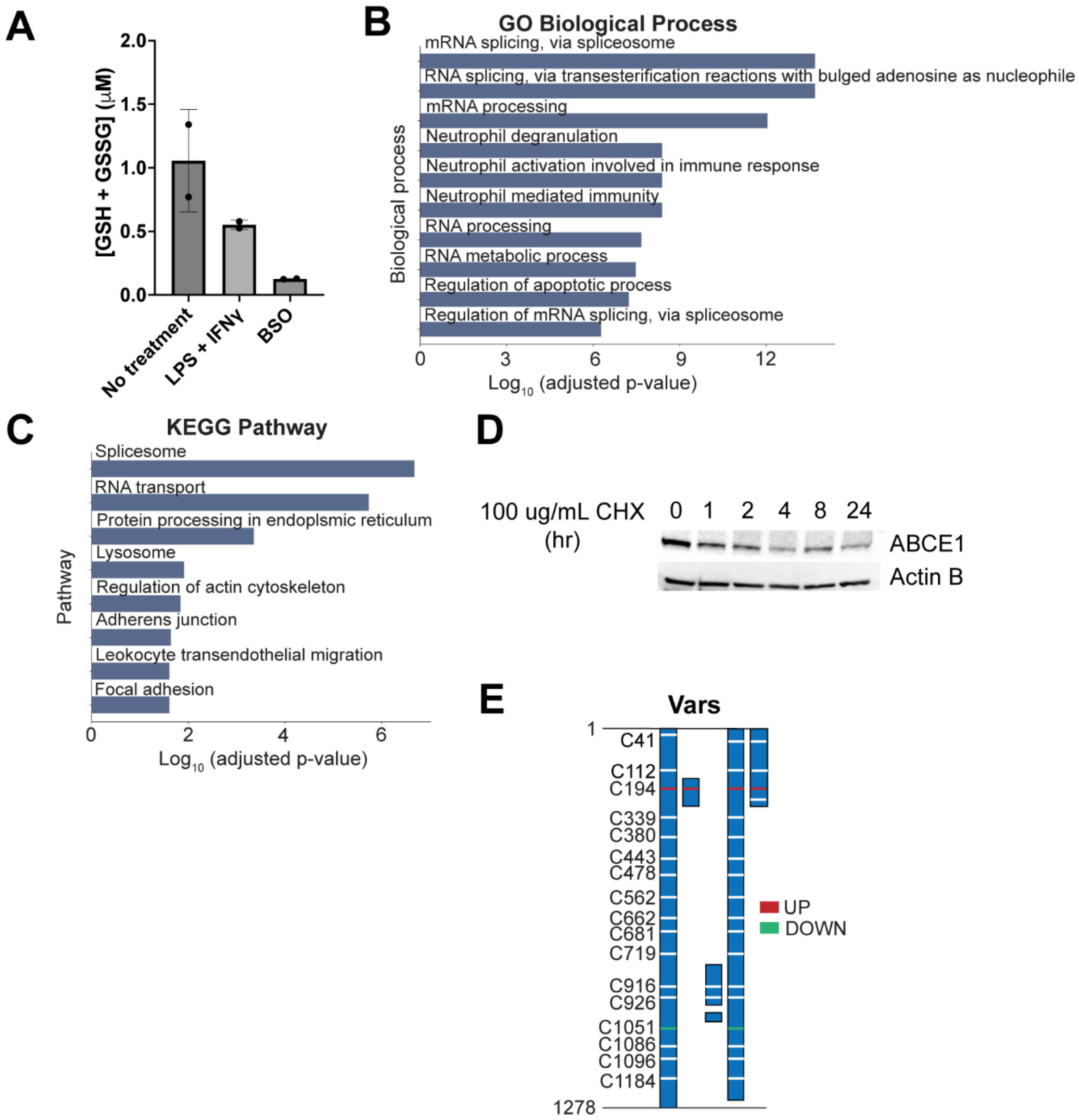


**Figure S14. A)** Amount of GSH detected in cells with or without LPS+IFNγ treatment. BSO was the negative control of the assay. **B)** GO biological process analysis of cysteines quantified with SP3-Rox that showed more reduced redox states upon LPS+IFNγ treatment. **C)** KEGG pathway analysis of cysteines quantified with SP3-Rox that showed more reduced redox states upon LPS+IFNγ treatment. **D)** Expression of ABCE1 upon treatment of translational inhibitor. **E)** Difference of redox states of cysteines quantified with SP3-Rox with or without LPS+IFNγ treatment in exemplary proteins with different splice forms. LPS+IFNγ treatment was 100 ng/mL LPS and 20 ng/mL IFNγ for 24 h at 37 °C. Experiments were performed in triplicates in iBMDM cells other than panel D, which was performed in HEK293T cells.

### Supplementary Tables

**Table S1-5.** Datasets corresponding to each figure, provided in the attached supplementary files.

**Table S6**. Plasmids used in this study.

| Name | Vector | Insert | Tag | Used for |
|---|---|---|---|---|
| p1 | pCDNA3 | Cytosol-TurboID | V5 (GKPIPNPLLGLDST) | Transient mammalian cytosol expression |
| p2 | pCDNA3 | ER-TurboID | V5 | Transient mammalian ER expression |
| p3 | pCDNA3 | Golgi-TurboID | 3xHA (YPYDVPDYA) | Transient mammalian Golgi expression |
| p4 | pCDNA3 | Mito-TurboID | V5 | Transient mammalian mitochondria expression |
| p5 | pCDNA3 | Nucleus-TurboID | 3xHA | Transient mammalian nucleus expression |
| p6 | FUGW | Cytosol-TurboID | V5, EGFP | Stable mammalian cytosol expression |
| p7 | FUGW | ER-TurboID | V5, EGFP | Stable mammalian ER expression |
| p8 | FUGW | Golgi-TurboID | 3xHA, EGFP | Stable mammalian Golgi expression |
| p9 | FUGW | Mito-TurboID | V5, EGFP | Stable mammalian mitochondria expression |
| p10 | FUGW | Nucleus-TurboID | 3xHA, EGFP | Stable mammalian nucleus expression |
| p11 | pVSVG | | | Producing lentivirus |
| p12 | Δ8.9 | | | Producing lentivirus |

**Table S7**. Antibodies used in this study.

| Name | Catalog number | Dilution (For western blot) |
|---|---|---|
| V5 Mouse mAb | Invitrogen R960-25 | 1:3000 |
| β-Actin Rabbit mAb | Abclonal AC038 | 1:10000 |
| Tomm 20 Rabbit mAb | Invitrogen MA5-32148 | 1:2000 |
| ABCE1 Rabbit mAb | Abclonal A9135 | 1:1000 |
| IRDye® 800CW Goat anti-Mouse IgG | LI-COR 92632210 | 1:5000 |
| IRDye® 800CW Goat anti-Rabbit IgG | LI-COR 92632211 | 1:5000 |
| IRDye® 800CW Streptavidin | LI-COR 92632230 | 1:5000 |

**Table S8.** Conditions of Liquid-chromatography (LC).

| Parameter | Condition |
|---|---|
| Column | 100 μM ID fused silica capillary packed in-house with bulk C18 reversed phase resin (particle size, 1.9 μm; pore size, 100 Å; Dr. Maisch GmbH) |
| Mobile phase | Buffer A: water with 3% DMSO and 0.1% formic acid<br>Buffer B: 80% acetonitrile with 3% DMSO and 0.1% formic acid |
| Gradient and flow rate | 0 – 5 min, 3 – 10% B, 300 nL/min<br>5 – 64 min, 10 - 50% B, 220 nL/min<br>64 – 70 min, 40 - 95% B, 250 nL/min |
| Run time | 70 minutes |
| Injection volume | 5 uL |

**Table S9.** Files in Proteomics Identification Database (PRIDE) datasets.

| Figure | File name | Experiment |
|---|---|---|
| 1 (HEK 293T) | 2020-10-05-KB-70min_FAIMS_3cv_-35_-45_-55_OTOT_SY41-1<br>2020-10-05-KB-70min_FAIMS_3cv_-35_-45_-55_OTOT_SY41-2 | Cys-Whole lysate |
| | 2022-08-25-KB-noFAIMS-SY87-1<br>2022-03-24-KB-SY87-1 | Cyto-Cys-LoC |
| | 2022-08-25-KB-noFAIMS-SY87-3<br>2022-03-24-KB-SY87-3 | ER-Cys-LoC |
| | 2022-08-25-KB-noFAIMS-SY87-4<br>2022-03-24-KB-SY87-4 | Golgi-Cys-LoC |
| | 2022-08-25-KB-noFAIMS-SY93-T500<br>2022-05-15-KB-nofaims-SY91-5a | Mito-Cys-LoC |
| | 2022-08-25-KB-noFAIMS-SY87-5<br>2022-03-24-KB-SY87-5 | Nuc-Cys-LoC |
| 2 (HEK 293T) | 2022-06-14-KB-NoFAIMS-standard<br>2022-05-14-KB-nofaims-standard | Whole lysate |
| | 2022-05-03-KB-SY91-ZT-1a<br>2022-05-03-KB-SY91-ZT-1b | Cyto-TurboID |
| | 2022-05-03-KB-SY91-ZT-3a<br>2022-05-03-KB-SY91-ZT-3b | ER-TurboID |
| | 2022-05-03-KB-SY91-ZT-4a<br>2022-05-03-KB-SY91-ZT-4b | Golgi-TurboID |
| | 2022-05-03-KB-SY91-ZT-5a<br>2022-05-03-KB-SY91-ZT-5b | Mito-TurboID |
| | 2022-05-03-KB-SY91-ZT-6a<br>2022-05-03-KB-SY91-ZT-bb | Nuc-TurboID |
| 3 (HEK 293T) | 2022-08-10-noFAIMS-SY98-1-1<br>2022-08-10-noFAIMS-SY98-1-2<br>2022-08-10-noFAIMS-SY98-1-3 | Ctrl<br>Mito-TurboID |
| | 2022-08-11-KB-noFAIMS-SY98-3-1<br>2022-08-11-KB-noFAIMS-SY98-3-2<br>2022-08-11-KB-noFAIMS-SY98-3-3 | Dia-FBS<br>Mito-TurboID |

| | | |
|---|---|---|
| | 2022-08-11-KB-noFAIMS-SY98-4-1<br>2022-08-11-KB-noFAIMS-SY98-4-2<br>2022-08-11-KB-noFAIMS-SY98-4-3 | CHX<br>Mito-TurboID |
| | 2022-08-08-noFAIMS-SY98-5-1<br>2022-08-08-noFAIMS-SY98-5-2<br>2022-08-08-noFAIMS-SY98-5-3 | Dia-FBS + CHX<br>Mito-TurboID |
| | 2022-08-20-KB-noFAIMS-SY99-1-1<br>2022-08-20-KB-noFAIMS-SY99-1-2<br>2022-08-20-KB-noFAIMS-SY99-1-3 | Ctrl<br>Mito-Cys-LoC |
| | 2022-08-20-KB-noFAIMS-SY99-2-1<br>2022-08-20-KB-noFAIMS-SY99-2-2<br>2022-08-20-KB-noFAIMS-SY99-2-3 | Dia-FBS + CHX<br>Mito-Cys-LoC |
| | 2022-08-20-KB-noFAIMS-SY99-5-1<br>2022-08-20-KB-noFAIMS-SY99-5-2<br>2022-08-20-KB-noFAIMS-SY99-5-3 | Dia-FBS + CHX<br>ER-Cys-LoC |
| | 2022-08-20-KB-noFAIMS-SY99-6-1<br>2022-08-20-KB-noFAIMS-SY99-6-2<br>2022-08-20-KB-noFAIMS-SY99-6-3 | Dia-FBS + CHX<br>Golgi-Cys-LoC |
| 4 (iBMDM) | 2022-10-07-KB-SY104-1-1<br>2022-10-07-KB-SY104-1-2<br>2022-10-07-KB-SY104-1-3 | Ctrl<br>Mito-Cys-LOx |
| | 2022-10-07-KB-SY104-2-1<br>2022-10-07-KB-SY104-2-2<br>2022-10-07-KB-SY104-2-3 | Dia-FBS + CHX<br>Mito-Cys-LOx |
| 5 (iBMDM) | 2022-10-07-KB-AJ-2stepredox-NT-1<br>2022-10-07-KB-AJ-2stepredox-NT-2<br>2022-10-07-KB-AJ-2stepredox-NT-3<br>2022-11-23-KB-AJ-2stepredox-NT-1-rerun<br>2022-11-23-KB-AJ-2stepredox-NT-2-rerun<br>2022-11-23-KB-AJ-2stepredox-NT-3-rerun | Ctrl<br>Mito-Cys-LOx |
| | 2022-10-07-KB-AJ-2stepredox-LPS-1<br>2022-10-07-KB-AJ-2stepredox-LPS-2<br>2022-10-07-KB-AJ-2stepredox-LPS-3<br>2022-11-23-KB-AJ-2stepredox-LPS-1-rerun<br>2022-11-23-KB-AJ-2stepredox-LPS-2-rerun<br>2022-11-23-KB-AJ-2stepredox-LPS-3-rerun | LPS<br>Mito-Cys-LOx |

| | | |
|---|---|---|
| | 2022-11-19-KB-noFAIMS-MV-SY-B16-mito-turboID-NT-2<br>2022-11-19-KB-noFAIMS-MV-SY-B16-mito-turboID-NT-4<br>2022-11-19-KB-noFAIMS-MV-SY-B16-mito-turboID-NT-6<br>2022-11-23-KB-noFAIMS-MV-SY-B16-mito-turboID-NT-2-rerun<br>2022-11-23-KB-noFAIMS-MV-SY-B16-mito-turboID-NT-4-rerun<br>2022-11-23-KB-noFAIMS-MV-SY-B16-mito-turboID-NT-6-rerun | Ctrl<br>SP3-Rox |
| | 2022-11-19-KB-noFAIMS-MV-SY-B16-mito-turboID-LPS-1<br>2022-11-19-KB-noFAIMS-MV-SY-B16-mito-turboID-LPS-3<br>2022-11-19-KB-noFAIMS-MV-SY-B16-mito-turboID-LPS-5<br>2022-11-23-KB-noFAIMS-MV-SY-B16-mito-turboID-LPS-1-rerun<br>2022-11-23-KB-noFAIMS-MV-SY-B16-mito-turboID-LPS-3-rerun<br>2022-11-23-KB-noFAIMS-MV-SY-B16-mito-turboID-LPS-5-rerun | LPS<br>SP3-Rox |

### Methods

**Cloning of different TurboID constructs.** List of plasmids with detailed information used in this study can be found in **Table S6**. PCR fragments of TurboID with different localization sequences were amplified using Phusion polymerase (Berkeley). The destination vectors and PCR products were both digested using standard enzymatic restriction and cleaned up, followed by T4 DNA ligation. Ligated products were transformed into TOP10 competent cells and sent for sequence to confirm the cloning.

**Cell culture.** Cell culture reagents including Dulbecco's phosphate-buffered saline (DPBS), Dulbecco's Modified Eagle Medium (DMEM) media, and penicillin/streptomycin (Pen/Strep) were purchased from Fisher Scientific. Fetal Bovine Serum (FBS) were purchased from Avantor Seradigm (lot # 214B17). All cell lines were obtained from ATCC and were maintained at a low passage number (< 20 passages). HEK293T (ATCC: CRL-3216) cells were cultured in DMEM supplemented with 10% FBS and 1% antibiotics (Penn/Strep, 100 U/mL). Immortalized bone marrow derived macrophages (iBMDMs) were generated by immortalizing murine BMDMs via overexpression of V-fraf and V-myc with J2 virus to generate immortalized BMDB (iBMDM) [1] iBMDMs were cultured in DMEM supplemented with 10% FBS, 1% antibiotics (Penn/Strep, 100 U/mL) and 5% (v/v) conditioned media containing macrophage colony stimulating factor (M-CSF) [2] produced by CMG cells to induce differentiation to BMDMs. Media was filtered (0.22 µm) prior to use. Cells were maintained in a humidified incubator at 37 °C with 5% CO2. Cell lines were tested for mycoplasma using the Mycoplasma Detection Kit (InvivoGen).

**Transfection of TurboID.** Cells were transfected at 70-80% confluency. For a 10-cm plate, plasmid (5 ug), serum-free DMEM (350 uL) and PEI (25 uL of 1 mg/mL) were mixed and incubated for 15 min at room temperature (RT), followed by adding dropwise to the cells.

**Generation of cell lines with stable TurboId expression.** For preparation of lentiviruses, HEK 293FT cells in 10 cm plates were transfected at ~80-90% confluency with lentiviral vector FUGW containing the gene of interest (10 ug; Addgene #14883)with the lentiviral packaging plasmids pVSVG (4 ug; Addgene #8454) and Δ8.9 (8 ug; Addgene #2221) and 66 uL of Turbo DNAfectin3000 (Lamda Biotech Inc.) in antibiotic-free media for 6 h. The DNAfectin-containing media was replaced with fresh antibiotic-free media and the cells were left to incubate for 48 hours for lentiviral generation. The media was collected and cells were allowed to incubate for another 24 hours in fresh media. After 24 h, the lentivirus-containing media was collected, and added to the previously harvested media. All collected lentivirus-containing media was stored at 4 °C. 1/3 volume of Lenti-X concentrator (Takara Bio, Cat# 631231) was added to the total harvested media and incubated 16 h at 4 C. The lentivirus was pelleted at 1500 g for 45 mins at 4 °C and resuspended in 500 uL plain DMEM and stored in 100 uL aliquots at -80 °C.

To generate the stable cell lines, cells were infected at 75 % confluency, passaged 5 times, and selected via flow cytometry for positive EGFP signal.

**Database Construction.** Subcellular location annotations from CellWhere Atlas (accessed 2208), Human Protein Atlas (HPA) version 21.1 and UniProtKB/Swiss-Prot (2208_release) were aggregated. Unique proteins were established using UniProt protein identifiers. CellWhere localization, HPA main location and UniProt subcellular location columns were mined for specific location keywords (ex. 'golgi'). Proteins containing these keywords are reported in **Table S1.**

**Biotinylation with TurboID.** For transiently expressed TurboID, biotin labeling was initiated 24 h after transfection. 100 mM biotin stock was made in dimethyl sulfoxide (DMSO). Biotin was directly added to cells at a final concentration of 500 μM and incubated for 1 h at 37 °C. After washing with cold DPBS for 3 times, cells were harvested by centrifugation (4,500 *g*, 5 min, 4 °C), washed twice with cold DPBS, lysed in RIPA buffer (Fisher, Cat# AAJ62885AE), and clarified by centrifuging (21,000 *g*, 10 min, 4 °C). Protein concentrations were determined using a BCA protein assay kit (Thermo Fisher, Cat# 23227) and the lysate diluted to the working concentrations indicated below.

**Gel and western blot.** Lysate was normalized to 2 mg/mL and separated on a 4-20% SDS-PAGE gel. Gels were transferred to nitrocellulose membrane (Bio-Rad) and blocked in 5% (w/v) milk in TBS-T (Tris-buffered saline, 0.1% Tween 20) for 1 h at RT. Membranes were incubated with primary antibodies overnight (14-16 h) at 4 °C then washed 3 times with TBS for 5 mins. Membranes were then incubated with secondary antibodies for 1 h at RT and washed 3 times with TBS. For blots assessing biotin signal, membranes were incubated with a streptavidin-fluorophore conjugate overnight at 4 °C. Membrane was imaged on Bio-Rad ChemiDoc. Antibodies used were listed in **Table S7**. ImageJ was used to normalize and quantify the band intensity.

**Proteomic sample preparation for TurboID, Cys-LoC and Cys-LOx.** Biotinylated lysates (500 μL of 1 mg/mL, prepared as described above) were labeled with either 2 mM IAA for Cys-LoC or 2 mM IPIAA-L for Cys-LOx for 1h at RT. 50 uL Pierce streptavidin agarose beads were washed with RIPA and incubated with lysates for 2 h at RT. The proteins bound to beads were washed once with 1 mL 2M urea in RIPA, twice with 1 mL RIPA, and 3 times with 1 mL PBS. The beads were resuspended in 200 μL 6 M urea, reduced with 1 mM DTT for 15 min at 65 °C, and labeled with 2 mM IAA for Cys-LoC or 2 mM IPIAA-H for Cys-LOx for 1h at RT. Then, beads were washed with PBS and resuspended in 200 μL 2 M urea. 3 μL of 1 mg/mL trypsin solution (Washington) was added. Proteins were digested off the bead overnight at 37 ºC with shaking. For one-step TurboID workflow, peptides were desalted with C18 column and analyzed by LC-MS/MS. For Cys-LoC and Cys-LOx, after digestion, CuAAC was performed with biotin-azide (4 μL of 200 mM stock in DMSO, final concentration = 4 mM), TCEP (4 μL of fresh 50 mM stock in water, final concentration = 1 mM), TBTA (12 μL of 1.7 mM stock in DMSO/*t*-butanol 1:4, final concentration = 100 μM), and CuSO4 (4 μL of 50 mM stock in water, final concentration = 1 mM) for 1h at RT. 20 μL Sera-Mag SpeedBeads Carboxyl Magnetic Beads, hydrophobic (GE Healthcare, 65152105050250, 50 μg/μL, total 1 mg) and 20 μL Sera-Mag SpeedBeads Carboxyl Magnetic Beads, hydrophilic (GE Healthcare, 45152105050250, 50 μg/μL, total 1 mg) were mixed and washed with water three times. The bead slurries were then transferred to the CuAAC samples, incubated for 5 min at RT with shaking (1000 rpm). Approximately 4 mL acetonitrile (> 95% of the final volume) was added to each sample and the mixtures were incubated for 10 min at RT with shaking (1000 rpm). The beads were then washed (3 × 1 mL acetonitrile) with a magnetic rack. Peptides were eluted from SP3 beads with 100 μL of 2% DMSO in MB water for 30 min at 37 °C with shaking (1000 rpm). The elution was repeated with 100 µL of 2% DMSO in MB water. For each sample, 50 µL of NeutrAvidin Agarose resin slurry (Pierce, 29200) was washed three times in 10 mL IAP buffer (50 mM MOPS pH 7.2, 10 mM sodium phosphate, and 50 mM NaCl buffer) and then resuspended in 800 µL IAP buffer. Peptide solutions eluted from SP3 beads were then transferred to the NeutrAvidin Agarose resin suspension, and the samples were rotated for 2 h at RT. After incubation, the beads were pelleted by centrifugation (21,000 *g*, 1 min) and washed (3 × 1 mL PBS, 6 × 1 mL water). Bound peptides were eluted twice with 60 µL of 80% acetonitrile in MB water containing 0.1% FA. The first 10 min incubation at RT and the second one at 72 °C. The combined eluants were dried (SpeedVac), then reconstituted with 5% acetonitrile and 1% FA in MB water and analyzed by LC-MS/MS.

**Crude mitochondria extraction.** HEK293T cells stably expressing mito-TurboID were plated in four 15-cm dishes. At 90% confluency, cells were harvested by centrifugation (4,500 *g*, 5 min, 4°C), washed twice with cold DPBS. The pellets were suspended in the MSHE buffer (70 mM sucrose, 210 mM mannitol, 5.0 mM HEPES, 1.0 mM EGTA) and homogenized in a glass grinder by 25 up-and-down passes of the pestle. The homogenate was then pelleted (1,100 *g*, 10 min, 4°C). The supernatant was transferred to a new tube and ultracentrifuged. (14,000 *g*, 10 min, 4°C). The supernatant was then saved as non-mito portion and the pellet was saved as mito portion.

**GC-MS for detecting biotin molecules inside cells.** HEK293T cells were plated in 15-cm dishes. At 90% confluency, 500 µM Biotin was added and incubated for 1 h at 37°C. Intact HEK293T cells or mitochondria isolated from HEK293T cells as described above were extracted and analyzed using GC/MS to quantify biotin levels. For extraction, 15 mL tubes containing 293T cells or isolated mitochondria were placed on ice. To remove residual biotin - containing culture medium, tissue culture cell pellets were first quickly washed with ice-cold 0.9% (w/v) NaCl. The cells or mitochondria pellets were immediately treated with 500 µL of ice-cold MeOH and 200 µL water containing 1 µg of the internal standard norvaline. Next, 500 µL of chloroform was added, after which samples were vortexed for 1 min and then spun at 10,000 *g* for 5 min at 4°C. The aqueous layer was transferred to a GC-MS sample vial and dried overnight using a refrigerated CentriVap. Once dry, samples were resuspended in 20 µL of 2% (w/v) methoxyamine in pyridine and incubated at 37°C for 45 min. This was followed by addition of 20 µL of MTBSTFA + 1% TBDMSCl (Ntert-Butyldimethylsilyl-N-methyltrifluoroacetamide with 1% tertButyldimethylchlorosilane), mixing, and incubation for an additional 45 min at 37°C. Samples were run as previously described [3], and analyzed using Agilent MassHunter software.

**Immunocytochemistry.** Mito-TurboID-EGFP stably expressed HEK293T cells were grown on glass coverslips in a 24-well plate. At 75% confluency, 500 µM Biotin was added and incubated for 1 h at 37°C. Cells were washed with PBS 3 times. Cells were fixed in 5% formalin in PBS for 15 min at RT and permeabilized by 0.1% Triton X-100 in PBS for 6 min at RT. Cells were blocked with 1% BSA for 1 hr at RT. For biotinylation detection, cells were treated with Streptavidin, Alexa Fluor™ 594 (1:500, Thermo Fisher, S11227) for 1 hr at RT. For mitochondrial localization, cells were treated with TMRE (1:1000, Invitrogen, T669) for 10 min at 37°C. Nucleus was detected with Dapi dye. Coverslips were mounted. Confocal images were obtained using Zeiss LSM 800

Confocal Laser Scanning Microscope. For localization analysis, the Coloc2 module in Image J was used with default settings.

**Generating streptavidin background dataset.** To generate the streptavidin background dataset, 2 individual negative control experiments for the TurboID workflow were prepared as previously described. In the negative control experiments, no TurboID fusion protein was expressed, and no exogenous biotin was added. Proteins were identified as "streptavidin background" if they were present in the streptavidin background dataset. The dataset contained a total of 966 proteins in aggregate.

**Respirometry.** Rates of oxygen consumption and extracellular acidification were measured using an Agilent Seahorse XFe96 Analyzer. Briefly, iBMDMs were plated at $7.5 \times 10^3$ per well in a 96-well Seahorse XF cell plate. After a 48-hour incubation, the cells were treated with 100 ng/mL lipopolysaccharide (LPS) and 20 ng/mL interferon gamma (IFNγ) for 24 hours. To mimic cysteine profiling experiments, 100 µg/mL cycloheximide was added to the cells for 6 hours prior to conducting respirometry experiments. At the time of experiment, iBMDM growth media was replaced with respirometry assay medium which consisted of unbuffered DMEM (Sigma #5030) supplemented with 2 mM pyruvate, 10 mM glucose, 2 mM glutamine, and 5 mM HEPES. Respiration was measured at baseline and in response to acute treatment with 2 µM oligomycin, FCCP (two sequential pulses of 0.75 µM), and 0.2 µM rotenone with 1 µM antimycin A. All respiratory parameters were calculated as previously described [4].

**RNA Isolation and qPCR analysis of iBMDMs.** Immortalized iBMDMs were plated at $1 \times 10^5$ per well in 12-well plates. After a 48-hour incubation, cells were treated with 100 ng/mL LPS and 20 ng/mL IFNγ for 24 hours. To maintain consistency with cysteine profiling experiments, 100 µg/mL cycloheximide was added to the cells for 6 hours prior to cell lysis. Following cell lysis, RNA was isolated using the RNeasy Mini kit (Qiagen). cDNA was synthesized with the high-capacity cDNA reverse transcription kit (Applied Biosystems). RT-qPCR assay was performed using PowerUp SYBR Green qPCR Master Mix kit (Applied Biosystems) on a QuantStudio 5 RT-PCR (Applied Biosystems). Relative gene expression values were calculated using the delta-delta Ct methods and Rplp0 was used as the control gene.

**Proteomic sample preparation for SP3-Rox of iBMDMs.** iBMDM cells were treated either with or without 100 ng/mL LPS and 20 ng/mL IFNγ for 24 h at 37 °C. Cells were then harvested and SP3-Rox procedure was carried out as reported [5].

**Liquid-chromatography tandem mass-spectrometry (LC-MS/MS) analysis.** The samples were analyzed by liquid chromatography tandem mass spectrometry using a Thermo Scientific™ Orbitrap Eclipse™ Tribrid™ mass spectrometer. Peptides were fractionated online using a 18 cm long, 100 µM inner diameter (ID) fused silica capillary packed in-house with bulk C18 reversed phase resin (particle size, 1.9 µm; pore size, 100 Å; Dr. Maisch GmbH). The 70-minute water-acetonitrile gradient was delivered using aThermo Scientific™ EASY-nLC™ 1200 system at different flow rates (Buffer A: water with 3% DMSO and 0.1% formic acid and Buffer B: 80% acetonitrile with 3% DMSO and 0.1% formic acid). The detailed gradient includes 0 – 5 min from 3 % to 10 % at 300 nL/min, 5 – 64 min from 10 % to 50 % at 220 nL/min, and 64 – 70 min from

50 % to 95 % at 250 nL/min buffer B in buffer A **(Table S8)**. Data was collected with charge exclusion (1, 8,>8). Data was acquired using a Data-Dependent Acquisition (DDA) method consisting of a full MS1 scan (Resolution = 120,000) followed by sequential MS2 scans (Resolution = 15,000) to utilize the remainder of the 1 second cycle time. Precursor isolation window was set as 1.6 and normalized collision energy were set as 30%. Details of MS data can be found in **Table S9**. All MS data was deposited to the ProteomeXchange Consortium via the PRIDE[6,7] partner repository with the dataset identifier PXD039626

**Protein, peptide, and cysteine identification.** Raw data collected by LC-MS/MS were searched with MSFragger (v3.3) and FragPipe (v19.0). The proteomic workflow and its collection of tools was set as default. Precursor and fragment mass tolerance was set as 20 ppm. Missed cleavages were allowed up to 1. Peptide length was set 7 - 50 and peptide mass range was set 500 - 5000. For Cys-LoC, Cysteine residues were searched with differential modification C+463.2366. For Cys-LOx and SP3-Rox, MS1 labeling quant was enabled with Light set as C+463.2366 and Heavy set as C+467.2529. MS1 intensity ratio of heavy and light labeled cysteine peptides were reported[8]. Calibrated and deisotoped spectrum files produced by FragPipe were retained and reused for this analysis. Custom python scripts were implemented to compile labeled peptide datasets. Unique proteins, unique cysteines, and unique peptides were quantified for each dataset. Unique proteins were established based on UniProt protein IDs. Unique peptides were found based on sequences containing a modified cysteine residue. Unique cysteines were classified by an identifier consisting of a UniProt protein ID and the amino acid number of the modified cysteine (ProteinID_C#); residue numbers were found by aligning the peptide sequence to the corresponding UniProt protein sequence. When there are multiple cysteines in one peptide, all the modified cysteine residue numbers will be reported as ProteinID_C#_C#..

**Data analysis.** For the subcellular annotation, our customized localization database was used to cross referenced with the proteins or cysteines identified. Proteins were counted as localized in the compartment that the TurboID fusion protein targeted even if they contained multiple localization annotations. For Cys-LOx, the medium of heavy to light ratios for the same cysteine residue from cysteine peptides of different charges and miss cleavages in the same sample was calculated. Means of reported logged ratio values for each condition (+/- LPS+IFNγ) were calculated for all replicates per condition, and the difference of the log2 mean values were reported. T-test was performed on the raw ratios to generate p-values. % oxidation for a cysteine was calculation based on heavy to light ratio via the following formula: (R/(1+R))*100, using unlogged ratios. When calculating oxidation difference, relative oxidation changes between two cellular conditions were reported by calculating the change of heavy to light ratios between treated and untreated samples.

**Splice variant analysis.** To obtain candidate proteins for splice variant analysis, we first filtered for proteins with at least one reduced and one oxidized cysteine. Fourteen proteins were identified, all of which were searched on ensembl genome browser and were found to have multiple protein coding splice splice variants. Three proteins were randomly selected as examples to compare cysteines present. For this subset, ensembl translation protein sequences for all variants were aligned using ensemble' multiple alignment tool, Clustal Omega. Alignments were exported and used to map Sp3-Rox-identified cysteines and their redox states.
